# Immune-profiling of T helper 1 (Th1), Th2 and Th17 signatures in murine splenocytes by targeting intracellular cytokines

**DOI:** 10.1101/2024.04.27.591473

**Authors:** Soumik Barman, Aisling Kelly, Danica Dong, Arsh Patel, Michael J. Buonopane, Jake Gonzales, Ben Janoschek, Andrew Draghi, David J. Dowling

## Abstract

Functional cytokines shape both innate and adaptive immune responses in the host after infection or immunization. Deep immunophenotyping of the key functional cytokine signatures associated with T cells in murine lymphoid tissue, especially in the spleen, is challenging. Using spectral flow cytometry, we developed a 17-parameter panel to profile major immune cell subsets along with T cells, memory phenotypes and functional cytokines in murine splenocytes in steady state as well as in stimulated conditions. This panel dissects the memory T cell compartment via CD62L and CD44 expression after mitogen stimulation. To profile T helper (Th) cells distribution after mitogen stimulation, established Th1 markers IFNγ, TNF and IL-2; Th2 markers IL-4/5 and the Th17 marker, IL-17, are included. This optimized multicolor spectral flow panel allows a detailed immune profiling of functional cytokines in the murine T cell compartment and might be useful for exploratory analysis of how these functional cytokines shape host immunity after infection or vaccination. Our panel could be easily modified, if researchers wish to tailor the panel to their specific needs.

## 1. PURPOSE AND APPROPRIATE SAMPLE TYPES

This 17-parameter panel was designed and optimized for the deep immunophenotyping and functional characterization of T cell subsets in murine spleen (Table 2 and Online Table 5). Functional cytokines (IFNγ, TNF and IL-2/4/5/17) in the T cell compartment initiate the innate immune responses after infection or immunization and establish a bridge between adaptive immune responses which ultimately confer long lasting protective immunity. Therefore, it is crucial to understand how functional cytokines influence the immune responses in the host. Our optimized panel allows profiling of Th1 cytokines (IFNγ, TNF and IL-2), Th2 cytokines (IL-4 and 5) and Th17 cytokine (IL-17) in effector CD4^+^ or CD8^+^ T cell compartment. Furthermore, the panel identifies B cells (B-1 and B-2), NK cells, gamma delta (γδ) T cells, NKT cells and γδNKT cells which are crucial in shaping host immunity. Importantly, to ensure broad applicability to the biomedical research field, we also validated the panel using splenocytes from two of the most predominantly utilized mice strains (C57BL/6 and BALB/c), in steady state along with phorbol myristate acetate (PMA) and ionomycin stimulated conditions *ex vivo*.

### 2. BACKGROUND

Despite the blooming of multi-dimensional immunophenotyping by spectral flow cytometry and the generation of over one hundred optimized multicolor immunofluorescence panels (OMIPs), there is currently no optimized high-dimensional panel established to target functional cytokines (Th1, Th2 and Th17 cells) in the murine model (Table 1). This 17-parameter spectral flow cytometry panel was developed to assess the functional cytokines production by conventional T cells which shapes the host immune responses upon natural infection or vaccinations (1). We previously demonstrated antigen specific responses or memory recall responses using microbial proteins (2,3) or peptides (4,5) and immunophenotyped functional cytokines in murine splenic T cell compartment after vaccination. Here, we expanded the flow cytometry panel using spectral flow technology by adding B cells, NK cells, and γδT cell markers, explored the memory T cell compartment and re-validated the panel upon mitogen stimulation using adult C57BL/6 and BALB/c mice strains (Figure 1, Table 2, Online Table 5 and Online Figure 8).

**Table 1.**
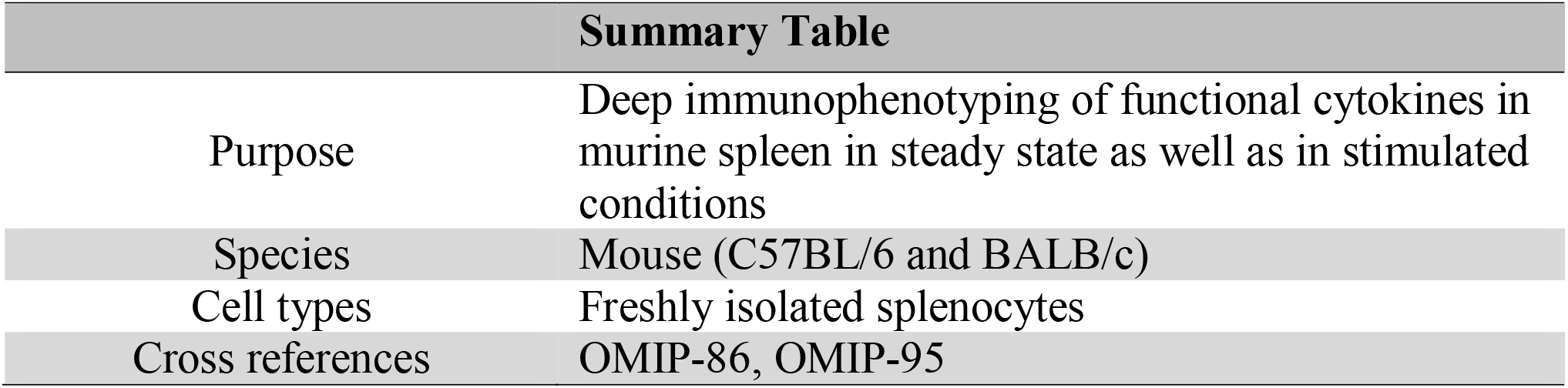
Summary table for the application of OMIP-XXX.

| Summary Table |  |
| --- | --- |
| Purpose | Deep immunophenotyping of functional cytokines in murine spleen in steady state as well as in stimulated conditions |
| Species | Mouse (C57BL/6 and BALB/c) |
| Cell types | Freshly isolated splenocytes |
| Cross references | OMIP-86, OMIP-95 |

**Table 2.**
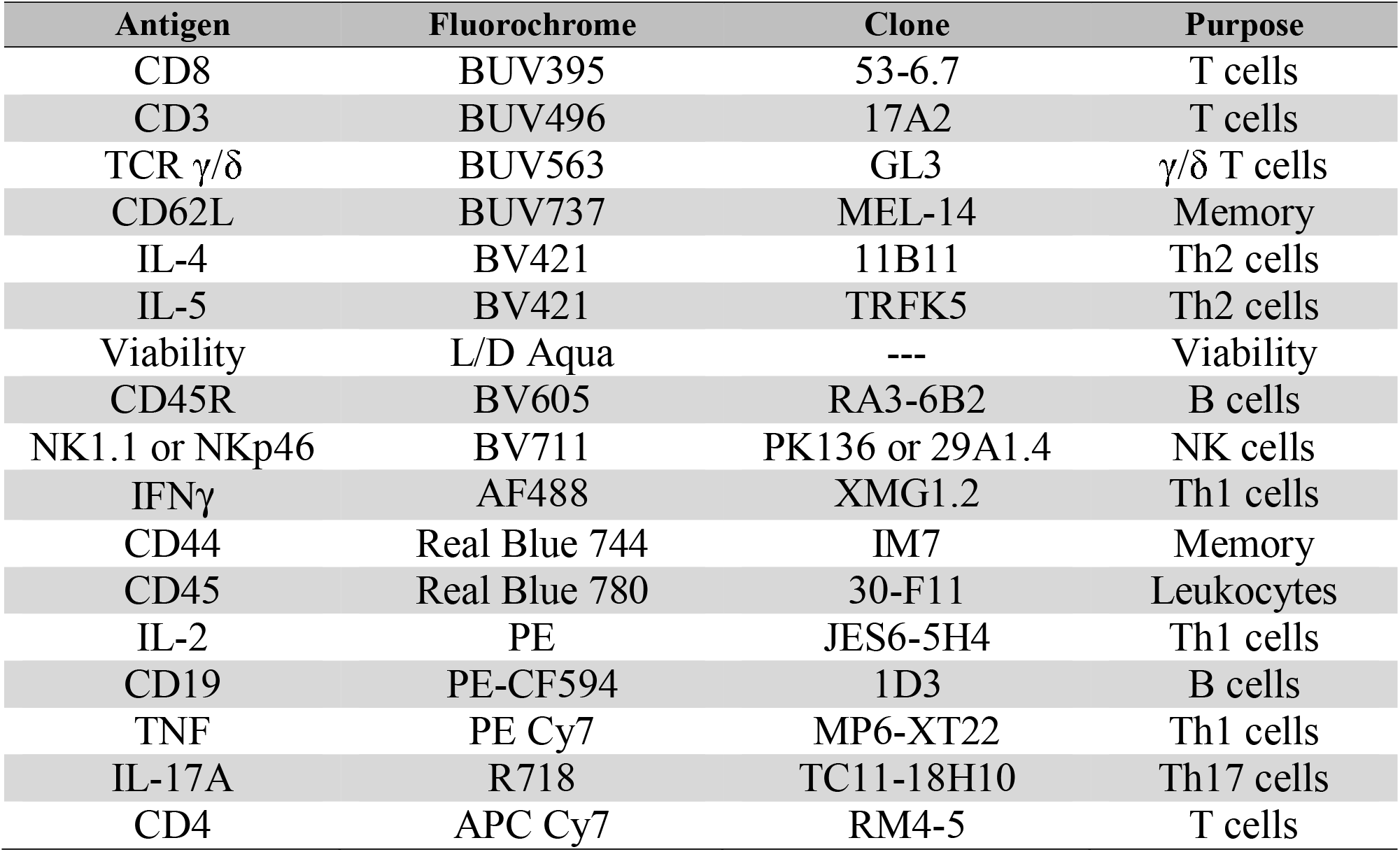
Panel and reagents for OMIP-XXX.

**Figure 1:**
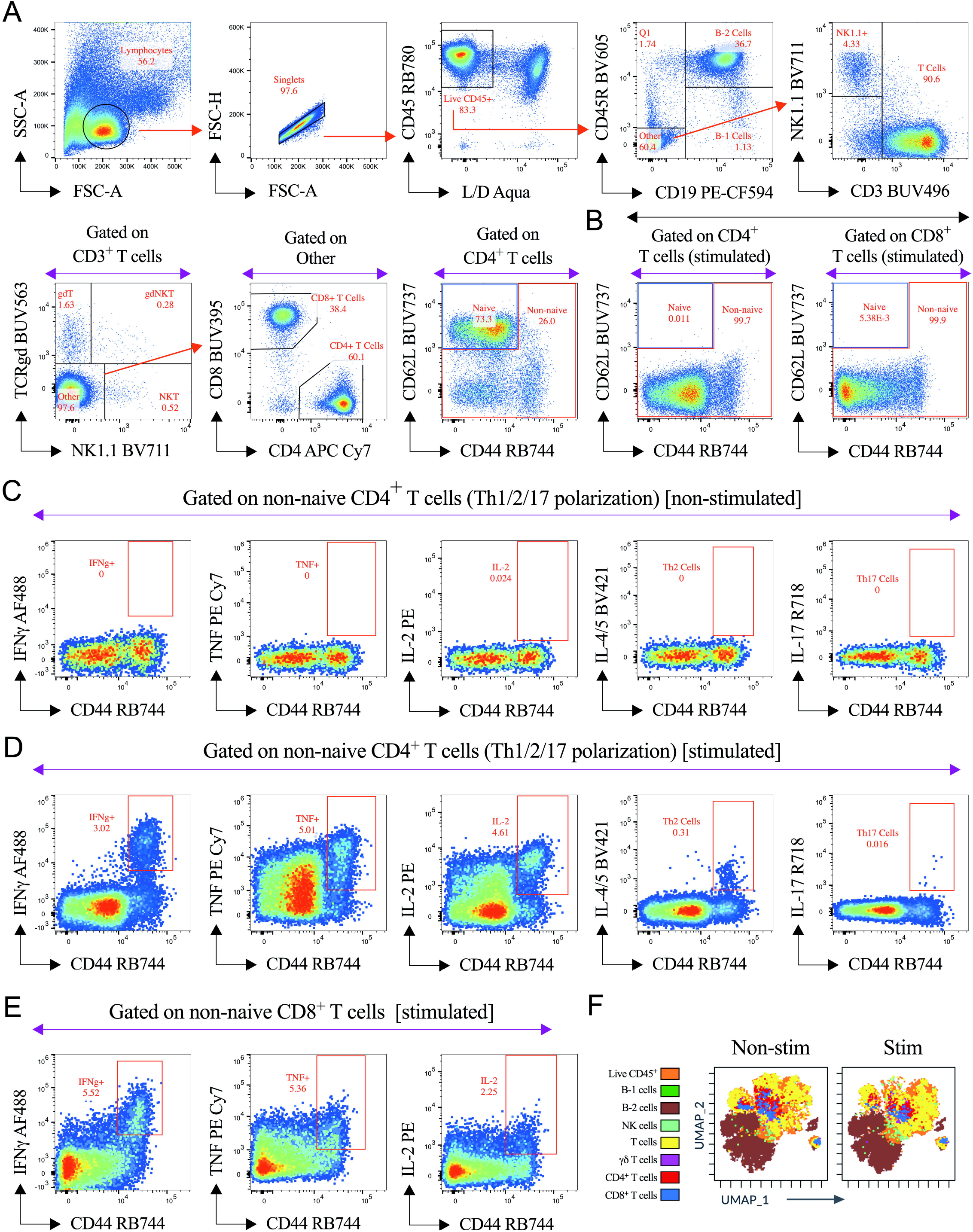
Representative gating strategy to profile functional cytokines and UMAP analysis for the visualization of the immune cell populations using 17-parameter OMIP-XXX in murine (C57BL/6) spleen. (A) Red arrows represent sequential gating for selected populations. After exclusion of cell debris and doublets, live splenic mononuclear cells were gated as CD45^+^ leukocytes. B cell populations (B-1 & B-2) were gated as CD19^+^ CD45R^+/-^. From the CD19^-^ CD45R^-^ subset, NK cells were phenotyped as CD3^-^ NK1.1^+^ subsets. Unconventional T cell populations were phenotyped in CD3^+^ T cell compartment according to the expression of TCRgd and NK1.1 and were portrayed as γδT cells (TCRgd^+^ NK1.1^-^), γδNKT cells (TCRgd^+^ NK1.1^+^) and NKT (TCRgd^-^NK1.1^+^) cells. After exclusion (TCRgd^-^NK1.1^-^) of unconventional T cells populations, conventional CD4^+^ and CD8^+^ T cell populations were targeted. Effector memory (CD44^+^ CD62L^-^), central memory (CD44^+^ CD62L^+^) along with double-negative cells (CD44^-^ CD62L^-^) portrayed as non-naive cells and excluded from naïve cell (CD44^-^ CD62L^+^) subsets in steady state as well as in (B) mitogen stimulated conditions for 6hrs. Th1 (CD4^+^ IFNγ^+^, CD4^+^ TNF^+^ and CD4^+^ IL-2^+^), Th2 (CD4^+^ IL-4/5^+^) and Th17 (CD4^+^ IL-17^+^) signatures were captured in murine splenic non-naïve T cell compartments in (C) steady state and in (D) 6hrs post-mitogen stimulations. (E) Th1 related cytokines (IFNγ, TNF and IL-2) were also detected in non-naïve compartments of CD8^+^ T cells. (F) The UMAP plots showing the phenotypic distribution of indicated cell populations. Indicated cell subsets were manually gated based on the phenotypic definition documented in (A). UMAP overlays via the Cytobank platform identified manually gated cellular phenotypes and displayed by the indicated color profiles in the cellular landscape either in steady state or in stimulated condition.

The white pulp in the murine spleen is enriched with lymphocytes, including B cells and T cells which orchestrate the host immune responses (6). We used CD45 as a common leukocyte marker and gated out dead cells using viability staining (Figure 1 and Online Figure 8). We phenotyped the B cell compartment and subdivided it according to CD19 and CD45R (B220) expression (7). B-1 cells (CD19^+^ CD45R^-^) and B-2 cells (CD19^+^ CD45R^+^) are two distinct types of B cells with different functions in the immune system. B-1 cells are involved in innate immunity and produce natural antibodies. B-2 cells, the more common type of B cells found in the spleen and lymph nodes, are a part of the adaptive immune system and generate specific antibodies in response to antigens (8,9). High-dimensionality (HD) reduction analysis using the UMAP algorithm also revealed the dominance of B-2 cells among the topography of live CD45^+^ cells (Figure 1F and Online Figure 8F).

NK cells are a type of lymphocytes that play a key role in the innate immune system. NK cells can recognize and kill cells that are infected with viruses or are transformed (such as cancer cells) without prior sensitization to specific antigens (10). We defined NK cells as CD45^+^ CD19^-^ CD45R^-^CD3^-^ NK1.1^+^ in C57BL/6 mice (Figure 1A) according to OMIP-95 (7). Anti-NK1.1 (PK136) antibody does not work in BALB/c mice strains because of an allelic discrepancy of the nkrp1b/c genes (11,12), which we also validated (Online Figure 7C) and used anti-NKp46 antibody (10,12) to target NK cells in BALB/c mice (Online Figures 7D and 8A).

Using NK1.1 or NKp46 along with TCRgd expression in CD3^+^ T cell populations, we identified unconventional NKT cells, γδT cells, and γδNKT cells in both C57BL/6 (Figure 1A) and BALB/c (Online Figure 8A) mice strains. Mainly NKT cells and γδT cells play a key role in the innate immune system despite there being reports of their involvement in triggering adaptive type memory responses through cytokine production (13-15). γδ T cells with an innate receptor (NK1.1 or NKp46), termed as γδNKT cells are believed to play a role in both innate and adaptive immunity, possibly in recognizing lipid antigens (16,17).

CD62L and CD44 markers are used to define naïve/memory phenotypes (Figure 1A, B and Online Figure 8A, B) in both CD4^+^ and CD8^+^ T cell compartments (7). Next, we focused on functional cytokine responses in the non-naïve/effector compartments of both CD4^+^ and CD8^+^ T cells after mitogen stimulation. A predominant portion of naïve CD4^+^ and CD8^+^ T cells become effector (Figure 1B and Online Figure 8B) after mitogen stimulation and successfully induced Th1 cells (CD4^+^ IFNγ^+^, CD4^+^ TNF^+^ and CD4^+^ IL-2^+^), Th2 cells (CD4^+^ IL-4/5^+^) and Th17 cells (CD4^+^ IL-17^+^) along with IFNγ, TNF and IL-2 induction in the effector CD8^+^ T compartment (Figure 1D, E and Online Figure 8D, E). We validated the consistency of the mitogen stimulated cytokine signatures across both C57BL/6 (Online Figure 9) and BALB/c (Online Figure 10) mice strains. In unstimulated conditions (only Brefeldin A treated) we did not find such signatures (Figure 1C and Online Figure 8C). Effector Th cells have important roles in host defenses. Th1 cells trigger macrophage-activation, whereas Th2 and Th17 cells are mainly involved in granulocyte recruitment and activation (18). Th1 cells predominantly provide defense against intracellular pathogens (18). Th2 cells have a potential role in clearing protozoan infections whereas Th17 cells act against extracellular bacteria and fungi (18). Th cells also participate in adaptive immunity after vaccination in an antigen-dependent manner by generating memory T cells that can deliver long-term protection (19).

In this OMIP, we optimized the staining panel for *ex vivo* analysis of functional cytokines secretion in CD4^+^ and CD8^+^ effector T cell compartments using murine splenocytes in steady state along with stimulated conditions. Panel optimization was completed using a Sony ID7000 spectral flow cytometer. Fluorochromes selection, modifications, and protocol details are documented in the online Supporting Text. All antibodies were individually titrated, and optimal concentrations were selected based on the lowest amount of antibody that provided the distinct separation between positive and negative populations with minimal background staining (Online Figures 4 and 7A,B). Fluorescence minus one (FMO) controls were used to define the gating strategy for all the surface markers (Online Figure 5). For the rare cytokine population, gating strategy was based on the Brefeldin A treated (unstimulated) controls (Figure 1C, Online Figures 6A-C and 8C). We observed APC-Cy7 tandem breakdown to its core tandem APC (Online Figure 11A), which has been previously observed by other groups (20,21). To overcome the phantom signal caused by the CD4 APC-Cy7 tandem breakdown, the APC channel was kept open (22). We utilized the Tandem Stabilizer (Online Table 4) and documented its effects up to 24h (Online Figure 11A). We did not experience TNF PE-Cy7 tandem breakdown to PE (Online Figure 6D-G) which is also a common pattern (21,22). We strongly recommend using FMO controls and Tandem Stabilizer to avoid these issues.

Taken together, this is the first spectral flow cytometry panel to profile major innate and adaptive immune cell populations in murine spleen along with intercellular Th1, Th2 and Th17 cells. Our optimized panel can also be easily expanded using additional markers. This panel could be employed to profile innate cytokine signatures after infection, or phenotype antigen specific functional cytokine profile by antigen stimulation (memory recall) of splenocytes using immunized animals.

## 3. SIMILARITY TO PUBLISHED OMIPS

This is the first OMIP available for immunophenotyping of murine splenic Th1, Th2 and Th17 cells based on intracellular cytokine signatures (Table 1, Table 2 and Online Table 5). To our knowledge, only few optimized panels focus on a broad range of intracellular cytokine secretion in murine models (OMIP-86). This OMIP exhibits minor overlap with OMIP-95 regarding the gating strategy to identify B cells, NKT cells, γδT cells, and γδNKT cells but none of those OMIPs targeted Th1, Th2 or Th17 cytokine signatures in mice.

## 4. STATEMENT OF ETHICAL APPROVAL OF ANIMAL EXPERIMENTS

All experiments were conducted following relevant institutional and national guidelines, regulations, approvals and in compliance with the ARRIVE (Animal Research: Reporting of *In Vivo* Experiments) guidelines. All experiments involving animals were approved by the Institutional Animal Care and Use Committee (IACUC) of Boston Children’s Hospital (protocol number 00001573). C57BL/6 and BALB/c mice were obtained from Jackson Laboratories (Bar Harbor, ME) and housed in specific pathogen-free conditions operated under the supervision of the Department of Animal Resources at Children’s Hospital (ARCH).

## AUTHOR CONTRIBUTIONS

**Soumik Barman:** Conceptualization; panel design; gating strategy; experimental work; data - collection, analysis, validation, visualization and deposition; writing - original draft; writing - review and editing. **Aisling Kelly & Danica Dong**: Murine spleen harvest and processing; writing - review and editing. **Arsh Patel**: Conceptualization; data - collection, analysis, validation and visualization; troubleshooting; software; writing □ review and editing. **Michael J. Buonopane**: Data - analysis, validation and visualization; troubleshooting; software; writing - review and editing. **Jake Gonzales**: Troubleshooting; writing - review and editing. **Ben Janoschek & Andrew Draghi II**: Conceptualization; panel design; data - analysis, validation and visualization; troubleshooting; software; writing - review and editing. **David J. Dowling**: Conceptualization; project administration; funding acquisition; mentorship and supervision; writing - review and editing.

## ACKNOWLEDGMENTS

The *Precision Vaccines Program* (PVP) is supported, in part, by the Boston Children’s Hospital, Department of Pediatrics. The authors thank the members of the PVP for helpful discussions, as well as Ofer Levy, Kevin Churchwell, Wendy Chung, Gary Fleisher, David Williams, and August Cervini for their support of the PVP. The authors thank Suzan Lazo, Director of the Cytometry Cores at Dana-Farber Cancer Institute (DFCI) for her support and feedback. Soumik Barman would like to thank Dr. Rajesh Panigrahi and Dr. Ravi Hingorani from the BD Life Sciences for their valuable advice on Real Blue fluorochromes during the panel optimization.

## FUNDING INFORMATION

The current study was supported by US National Institutes of Health (NIH)/National Institutes of Allergy and Infectious Diseases (NIAID) awards, including Adjuvant Discovery (75N93019C00044), Adjuvant Development (272201800047C) and Development of Vaccines for the Treatment of Opioid Use Disorder (272201800047C-P00003-9999) Program contracts (to David J. Dowling).

## CONFLICT OF INTEREST STATEMENT

Ben Janoschek and Andrew Draghi II are employees of Sony Biotechnology Inc., the manufacturer of the Sony ID7000 flow cytometer used in this study. Jake Gonzales is an employee of BioLegend Inc. David J. Dowling is a named inventors on patents relating to small molecule adjuvants assigned to Boston Children’s Hospital, is on the scientific advisory board of EdJen BioTech and serves as a consultant with Merck Research Laboratories/Merck Sharp & Dohme Corp. (a subsidiary of Merck & Co., Inc.). David J. Dowling is cofounder of Ovax Inc. The other authors declare no conflict of interest.

## DATA AVAILABILITY STATEMENT

Data are available in the main article, figure, tables and in the supporting information. Deposited raw fully stained quality-controlled data (FCS files) can be accessed through the Flow Repository (FR-FCM-Z7CH). The deposited data will be publicly available after the paper is published. Additional data which support the optimization of the study (i.e. FMOs, stained samples without tandem stabilizer) are available from the corresponding authors upon reasonable request.

## Notes

http://flowrepository.org/id/FR-FCM-Z7CH

