## Supporting Information for "Immune-profiling of T helper 1 (Th1), Th2 and Th17 signatures in murine splenocytes by targeting intracellular cytokines"

### Table of Contents


| Supporting Text | | Page |
|---|---|---|
| | Instrument (Sony ID7000) configuration, fluorochromes selection, modifications and protocol details are documented here. | S-3 to S-9 |

| Supplementary Tables | | Page |
|---|---|---|
| Online Table 1 | Biological samples, reagents, and algorithms. | S-10 |
| Online Table 2 | Reagents used to prepare 2X T Cell Media (TCM). | S-11 |
| Online Table 3 | Reagents used to prepare FACS Buffer. | S-11 |
| Online Table 4 | Additional reagents for flow cytometry assay. | S-11 |
| Online Table 5 | Table for flow antibodies used in the optimized 17-parameter panel. | S-12 |

| Supplementary Figures | | Page |
|---|---|---|
| Online Figure 1 | Graphical representation of the optical layout and configuration of the detectors on the Sony ID7000 spectral cell analyzer. | S-13 |
| Online Figure 2 | Similarity Index Matrix (SIM) of the optimized 17-color panel on the Sony ID7000 spectral cell analyzer. | S-14 |
| Online Figure 3 | Spillover Spreading Matrix (SSM) of the optimized 17-color panel on the Sony ID7000 spectral cell analyzer. | S-15 |
| Online Figure 4 | Optimal concentration of antibodies is defined by titrations. | S-16 |
| Online Figure 5 | Defining gating strategy using FMO controls. | S-18 |
| Online Figure 6 | Defining the gating strategy for cytokine$^+$ cells in CD8$^+$ T cell compartment and assessing of PE-Cy7 tandem breakdown. | S-19 |
| Online Figure 7 | Marker for NK cells iterations during panel optimization using C57BL/6 and BALB/c mice. | S-20 |
| Online Figure 8 | Panel compatibility in BALB/c mice. | S-21 |
| Online Figure 9 | Consistency of the cytokine signatures (Th1, Th2 & Th17) across C57BL/6 mice strain. | S-23 |
| Online Figure 10 | Consistency of the cytokine signatures (Th1, Th2 & Th17) across BALB/c mice strain. | S-25 |
| Online Figure 11 | Reasoning behind using the Tandem Stabilizer. | S-27 |
| Online Figure 12 | Dimensionality reduction analysis using the UMAP algorithm and distribution of surface markers in the cellular topography. | S-28 |
| Online Figure 13 | Dimensionality reduction analysis using UMAP algorithm and distribution of intracellular cytokine markers in the cellular topography. | S-29 |

**Supporting Text:**

#### Instrument configuration

This OMIP was developed using Sony ID7000 spectral cell analyzer equipped with 5 lasers (355nm, 405nm, 488nm, 561nm and 637nm) and 147 fluorescence detectors. The optical layout and fluorescence detectors configuration were documented in **Online Figure 1**. The UV (355nm) and Violet (405nm) lasers each have 3 individual PMTs on the lower range of the spectrum as well as a 32 channel PMT deck. The Blue (488nm) laser gives FSC and SSC measurements as well as has its own 32 channel PMT deck. The Yellow-Green (561nm) and Red (637nm) lasers have 26 and 19 detectors respectively in each of their detector decks. There is a diffraction grating system in front of each of the detector decks to distribute light evenly across the spectrum and enables ~10nm resolution from 494nm to 844nm.

Detector voltage setting was fine-tuned to provide optimal resolution for each channel according to Perfetto *et al* (1). FSC and SSC voltages were only manually adjusted to fit the lymphocytes populations. Quality control of the ID7000 is performed daily by the DFCI cytometry core using AlignCheck Particles (Sony, Cat# AE700510).

#### Strategy for the panel development

The goal of developing this OMIP was to generate a panel to phenotype murine intracellular cytokines (Th1, Th2 and Th17) in stimulated conditions. In the past, we, *Precision Vaccines Program*, phenotyped flu (2) or pertussis protein (3) specific and SARS-CoV-2 peptide specific (4,5) memory recall responses after vaccination or adjuvanted vaccination to understand the impact of certain adjuvants in the murine model by conventional flow cytometry. Here, we took the advantage of spectral flow cytometry and advanced our original conventional flow panel backbone by further adding B cells, NK cells, γδT cells along with memory markers using ID7000 spectral cell analyzer. All the markers included in the final version of the panel are shown in **Table 2** and in **Online Table 5**. Below are the details of the rationale for each fluorochrome specific marker selection.

CD8 (T cell marker) was assigned to BUV395 (a dim/moderate fluorophore in the UV spectrum) due to its unique spectral signature and less or negligible spectral spread on other fluorochromes according to previous experiences (2-5), which is also confirmed using SIM (**Online Figure 2**) followed by SSM (**Online Figure 3**).

CD3 (T cell marker) was assigned to BUV496 which is a dim/moderate fluorophore in the UV spectrum. BUV496 has spectral spread (SSM score 39.66, **Online Figure 3**) in Live/Dead Fixable Aqua because of the high similarity index (Similarity score: 0.64, **Online Figure 2**) between the fluorochrome pairs, but we have not faced a problem as Live/Dead Fixable Aqua positive cells were considered as the dead cells and gated out from the downstream analysis (**Figure 1** and **Online Figure 8**). Live/Dead Fixable Aqua was selected as viability dye according to previous experiences (2-5).

During the panel development, we tried to assign bright dyes to target dim markers. TCRgd (γδ T cell marker which is dim in expression) was assigned to a bright fluorophore BUV563. Less or negligible spectral spread on other fluorochromes was confirmed using SIM (**Online Figure 2**) followed by SSM (**Online Figure 3**).

Memory phenotype defining markers CD62L and CD44 were assigned to BUV737 and Real Blue (RB) 744 respectively. Although we have extensive experiences with CD44 PerCP-Cy5.5 (2-5), we decided to use CD44 RB744 for this spectral panel because of its unique spectral signature and negligible spectral spread on CD62L BUV737 and other fluorochromes as described in SIM (**Online Figure 2**) and SSM (**Online Figure 3**).

BV421 and BV711 are the bright fluorochromes in the violet spectrum. BV421 was assigned to detect intracellular cytokines IL-4 and IL-5 according to the previous experience (3). NK1.1 (in C57BL/6) and NKp46 (in BALB/c) were assigned to BV711 and validated (**Online Figure 7**) using two separate panels (**Figure 1** for C57BL/6 and **Online Figure 8** for BALB/c).

Due to the low level of expression (dim), B cell markers CD45R and CD19 were assigned to bright fluorophores BV605 and PE-CF594, respectively. Highly expressed leukocytes marker CD45 was assigned to a bright fluorophore RB780 in the blue spectrum, despite we did not see any drawback according to the SIM (**Online Figure 2**) followed by SSM (**Online Figure 3**).

Th1 cytokines IFNγ, IL-2 and TNF were assigned to AF488, PE and PE-Cy7 respectively, according to the previous experiences (2-5). To track down Th17 cytokine IL-17 (dim marker), we used a bright fluorophore R718 in the red spectrum.

For CD4 (T cell marker), we choose a dim fluorophore APC Cy7 in the red spectrum because of the high abundance of CD4 expression in murine splenic cells. We were aware of the tandem breakdown to its core tandem APC and therefore kept the APC channel blank (**Online Figure 11**). We used the Tandem Stabilizer (**Online Table 4**) to prevent the APC Cy7 tandem breakdown (**Online Figure 11A**). The Tandem Stabilizer presents the potential to add further markers in the APC channel without any interference.

**Panel optimization and titrations of antibodies**

After the selection of fluorophores either by previous experiences (2-5) or using SIM (**Online Figure 2**), next we titrated selected antibodies using freshly isolated and pulled murine (C57BL/6) splenic cells at an average of 2 million cells per test (per well). All antibodies were titrated in a two-fold dilution series except Live/Dead Fixable Aqua (**Online Figure 4**). We started with 1:20 dilution (5μL of test antibody in 100μL of buffer containing 2 million cells per well) for all the targeted antibodies and titrated up to 1:320 dilution (**Online Figure 4**). For Live/Dead Fixable Aqua, the starting dilution (using PBS) was 1:125 and titrated up to 1:1000 in a two-fold dilution series, except the inclusion of 1:800 dilution (**Online Figure 4**).

NK1.1 BV711 and NKp46 BV711 were titrated using splenic cells from C57BL/6 and BALB/c mice respectively (**Online Figure 7**). We also validated the previous report of allelic discrepancy of the nkrp1b/c genes in BALB/c mice strains (6,7). As a result, NK1.1 BV711 did not work in BALB/c mice strains (**Online Figure 7C**). We kept the panel same for BALB/c mice with alteration of NK1.1 BV711 to NKp46 BV711 (**Online Figure 8** and **Online Table 5**).

To titrate functional intracellular cytokines, we stimulated freshly isolated and pulled murine splenic cells using the Cell Activation Cocktail or mitogen (**Online Table 1**). Brefeldin A was used to block the cytokines. The details of the stimulation and staining procedures are available in the "**Mitogen stimulation**", "**Viability Stain and antibody preparation**", "**Surface antibody staining**" and "**Intracellular cytokine staining**" sections.

**Reference control**

The SSM should be similar despite the computation being done using cellular controls or antibody-capture beads (8). As we are dealing with rare populations of functional cytokines, it is challenging to ensure a positive signal using cellular controls. Therefore, we decided to use UltraComp eBeads™ Plus (**Online Table 4**) for computing both the SIM and SSM along with performing unmixing in this publication. The advantage of using beads is that it promises a positive or bright signal of the targeted antibody. Probably, beads are superior for computing SSM (9).

Optimal concentration (**Online Figure 4** and **Online Table 5**) of the selected antibody conjugates (per well) was plated in a 96 well round (U) bottom plate (**Online Table 1**) containing 75 µL of FACS buffer (**Online Table 3**). After thoroughly vortexing the UltraComp eBeads™ Plus, one drop of the bead was added in each well. For Live/Dead Fixable Aqua, one drop of ArC™ reactive beads (thoroughly vortexed) were added in a separate dry well, rested for 5 mins and 3 µL of the test antibody was added to it directly. After 20 mins of incubation at RT in dark condition, the samples were centrifuged (2200 RPM, 4 mins), resuspended and stored in PBS (200µL/well) at 4°C (in dark condition) until acquisition. For Live/Dead Fixable Aqua, after washing the beads were suspended in 140 µL of PBS followed by the addition of a drop of ArC™ negative beads (thoroughly vortexed). All the acquisitions in this publication were completed within 18h of the completion of stains with freshly prepared reference controls for each experiment unless otherwise mentioned. During acquisition in the ID7000 spectral cell analyzer, we acquired 10,000 reference beads per sample on every occasion, performed the unmixing using Weighted Least Squares Method (WLSM) as the unmixing algorithm and defined the SSM (**Online Figure 3**).

**Preparation of single cell suspension from murine spleen**

Spleens from 10-12 weeks old mice (C57BL/6 or BALB/c) were harvested in RPMI 1640 media containing 10% heat inactivated FBS (**Online Table 1**), aseptically. Harvested spleens were mechanically dissociated (gently) with the back of a 3mL syringe (**Online Table 1**) plunger and filtered through a pre-wetted 70 µm cell strainer (**Online Table 1**) in a 50mL centrifuge tube, aseptically. After centrifugation (1200 RPM, 10 min, RT), cells were treated with 1 mL ACK lysis buffer (**Online Table 1**) for 2 minutes at RT. Cells were washed immediately with RPMI 1640 media, passed through a 70 µm cell strainer again, and suspended in 3mL of RPMI 1640 media (supplemented with 10% heat inactivated FBS) in a 15mL centrifuge tube (**Online Table 1**). Cell concentrations were determined by Nexcelom Cellometer K2 using ViaStain™ AOPI staining solution and Cellometer counting chambers (**Online Table 1**). Splenocytes were plated at a density of up to 2 × $10^6$ cells/well in a 96 well round (U) bottom plate containing 180µL of 1X TCM (**Online Table 2**) and rested for 2 hours to restore the basal metabolic activity. To titrate intracellular antibodies, splenocytes were plated at a density of up to 6 × $10^6$ cells/well in a Costar® 48 well Flat Bottom Plate (**Online Table 1**) containing 540µL of 1X TCM and rested for 2 hours. Resting and stimulation were done using a humidified incubator at 37°C in the presence of 5% $CO_2$.

**Mitogen stimulation**

After resting, cells were stimulated for 6h using the Cell Activation Cocktail (mitogen) along with Brefeldin A (**Online Table 1**), which blocks the cytokine production and facilitates the optimal detection of the intracellular cytokines during spectral flow cytometry analysis. Either 20µL (for wells containing 2 × $10^6$ cells in 180µL of 1X TCM) or 60µL (for wells containing 6 × $10^6$ cells in 540µL of 1X TCM) of stimulation cocktail was added per well to achieve a total volume of either 200 or 600µL (per well) respectively. The concentration of the Cell Activation Cocktail (1:500) and Brefeldin A (1:1000) were determined according to our previous studies (2,3). After completion of stimulation, plates were kept at 4°C.

**Viability Stain and antibody preparation**

Next morning (within 14h), a cocktail of surface antibodies (**Online Table 5**) was prepared using FACS buffer (90µL/test) along with Brilliant Stain Buffer Plus (10µL/test, **Online Table 4**). The cocktail of ICS antibodies (**Online Table 5**) was prepared using 1X Perm/Wash buffer (90µL/test) along with Brilliant Stain Buffer Plus (10µL/test). Experimental fluorescence minus one (FMO) for either surface or intracellular antibodies were also made in a similar fashion.

Aqua Live/Dead staining solution (1:500 dilution in PBS) was prepared freshly during the staining procedure. For viability staining, cells were centrifuged (2200 RPM, 3 mins, 18°C), washed once with PBS (200µL/well) and treated with 100µL of Mouse Fc Block (**Online Table 4**) for 10 mins at RT. Next, cells were centrifuged and stained with Aqua Live/Dead stain (50µL/well) for 15 minutes at RT.

For titration of intracellular cytokines, at first mitogen-stimulated cells were harvested from 48 well flat bottom plate in a 15mL centrifuge tube. The attached cells at the bottom of the surface were recovered by adding 200µL of 0.5mM EDTA (**Online Table 1**) followed by a short incubation for 10min at RT and added to the master centrifuge tube. After centrifugation (1800 RPM, 3min, RT), cells were dissolved in 1 mL RPMI 1640, counted using Nexcelom Cellometer K2 and transferred to 96 well round (U) bottom plate as 2 × $10^6$ cells/well. After transfer, cells were washed and treated with FC blocker followed by viability staining procedure as described above.

For titration of surface antibodies, freshly isolated 2 × $10^6$ cells were plated per well, rested and directly subjected to viability staining.

**Surface antibody staining**

After viability stains, cells were washed (2200 RPM, 3 mins, 18°C) with PBS twice, and incubated with 100μL (per well) of cocktail of surface antibodies for the full panel run. For titration (surface antibodies), each well was incubated with 100μL of diluted single fluorophores in 100μL of FACS buffer. For titration of intracellular cytokines, cells were treated with 100μL of FACS buffer only. Co-staining with CD3 was performed for pre-gating of $CD3^{+/-}$ cells (**Online Figures 4** and **7A,B**) to evaluate certain antibodies if the biological expression pattern is moderate or rare. All the plates were incubated at 4°C for 30 minutes in the dark.

Next, after washing (2200 RPM, 3 mins, 4°C) twice with PBS, the wells (designated for surface antibodies titration only) were subjected to 1% paraformaldehyde (**Online Table 4**) for 20 minutes at 4°C, washed twice with PBS again, and stored in PBS at 4°C until acquisition.

**Intracellular cytokine staining**

After surface stain, cells were washed (2200 RPM, 3 mins, 4°C) twice with PBS. Next, we fixed and permeabilized the cells using 100μL (per well) of BD Cytofix buffer (**Online Table 4**) for 20 mins at 4°C in the dark. After fixation and permeabilization, cells were washed twice with 100μL of BD 1X Perm/Wash buffer and were subjected to intracellular staining (30 minutes at 4°C) using a cocktail (100μL/well) of targeted intracellular cytokines (**Online Table 5**). For the titration of intracellular cytokine antibodies, cells were treated with 100μL of diluted single fluorophores in 100μL of BD 1X perm/Wash buffer only and incubated at 4°C for 30 minutes in the dark.

After completion of the incubation, cells were washed twice using 1X perm/Wash buffer. Finally, cells were fixed in 1% paraformaldehyde for 20 minutes at 4°C, washed twice with PBS and stored in PBS (200μL/well) at 4°C until acquisition.

In certain conditions, cells were washed only once with PBS after fixation in 1% paraformaldehyde and subjected to additional steps for preservation to bypass the tandem breakdown.

**Preservation of the stained cells**

After fixation using 1% paraformaldehyde and washing once with PBS, cells were subjected to another washing step with PBS (200μL/well) containing Tandem Stabilizer (**Online Table 4**). After the last wash, finally cells were preserved in 200μL of PBS containing the Tandem Stabilizer at 4°C in the dark until acquisition.

**Data acquisition and analysis**

Within 18h of the completion of the staining procedures, data were acquired unless mentioned otherwise. After running single color reference beads and defining unmixing, test samples were analyzed. For the full panel run, a stop gate was applied on live $CD45^{+}$ cells with a count of 100K cells. After completion of the acquisition, all the fluorochrome combinations (NxN plots) were scrutinized using the “unmixing viewer”. If necessary, the distribution of the targeted populations was adjusted by activating the adjuster followed by dragging the populations and applied the new unmixing to the entire data set.

FCS files generated by ID7000 spectral cell analyzer, were analyzed using FlowJo 10.10 for Mac Operating System. Fluorescence minus one controls (FMOs) were used to justify the gating strategy for all the surface makers (**Online Figure 5**). For the rare cytokine population, gating strategy was based on the unstimulated (Brefeldin A treated) controls (**Figure 1C**, **Online Figure 6A-C** and **Online Figure 8C**). The gating strategy for the final panel using C57BL/6 strain was demonstrated in **Figure 1**. The panel was revalidated using BALB/c strain and demonstrated in **Online Figure 8**. **Online Figures 9** and **10** demonstrated the consistency of the panel to detect Th1, Th2 and Th17 signatures in C57BL/6 and BALB/c mice respectively after mitogen stimulation. UMAP analysis was performed using the Cytobank platform (**Online Figures 12** and **13**) where 23K events of live $CD45^+$ populations were sampled equally. For the UMAP run, the setting of neighbors and distance parameters were adopted from OMIP-95 (10). The parameters for UMAP run were Num neighbors= 80, Minimum distance = 0.7 and collapse outliers: Yes. By UMAP embedding, we overlaid manually gated major innate and adaptive immune cell lineages in the cellular landscape of live $CD45^+$ subset (**Figure 1F** and **Online Figure 8F**). Dimensionality reduction analysis identified the unique cellar landscape expressing the targeted surface markers (**Online Figure 12**) along with intracellular markers (**Online Figure 13**) assigned in the Z-axis channel.

**Online Table 1.** Biological samples, reagents, and algorithms.

| Reagent or Resource | Source | Identifier |
|---|---|---|
| Biological Samples | | |
| Splenocytes from 10-12 weeks old C57BL/6 and BALB/c mice were collected aseptically for the entire study. | All experiments involving animals were approved by the Institutional Animal Care and Use Committees (IACUC) of the BCH. | --- |
| Chemicals, Materials, Stimulants | | |
| Fetal Bovine Serum (FBS) | Gibco | 16000-044 |
| 70 µm Cell Strainer | Falcon, Corning | 352350 |
| 3mL syringe | BD | 309657 |
| ACK Lysis Buffer | Gibco | A10492-01 |
| ViaStain™ AOPI Staining Solution | Nexcelom | CS2-0106-5mL |
| Cellometer Counting Chambers | Nexcelom | CHT4-SD100-002 |
| Cell Activation Cocktail | BioLegend | 423301 |
| Brefeldin A Solution | BioLegend | 420601 |
| UltraPure™ 0.5M EDTA | Invitrogen | 15575-038 |
| 96 Well Round (U) Bottom Plate | ThermoFisher | 163320 |
| Costar® 48 well Flat Bottom Plate | Corning | 3548 |
| 15 mL Centrifuge Tube | CELLTREAT | 229411 |
| 50 mL Centrifuge Tube | CELLTREAT | 229421 |
| Software and Algorithms | | |
| FlowJo (10.10) for macOS | BD | https://www.flowjo.com/solutions/flowjo/downloads |
| UMAP | Cytobank | https://premium.cytobank.org/cytobank/ |
| Prism 10 (10.2.2) for macOS | GraphPad Software | https://www.graphpad.com |
| EndNote 21 | Clarivate | https://endnote.com |
| Microsoft 365 | Microsoft | https://www.microsoft.com/en-us/microsoft-365 |
| Deposited Data | | |
| FCS files for OMIP-XXX | FLOW Repository | http://flowrepository.org<br>Repository ID: FR-FCM-Z7CH |

**Online Table 2.** Reagents used to prepare 2X T Cell Media (TCM). After preparation and mixing the reagents aseptically, the media was passed through a 0.2 μM syringe filter (Corning, Cat# 431229) using a 50 mL syringe (BD, Cat# 309654) and kept at 4°C for storage. TCM was diluted to 1X using RPMI before usage.

| Reagent | Vendor | Identifier | Working concentration (in 1X) | mL/per 50 mL (in 2X) |
|---|---|---|---|---|
| FBS-HyClone | Cytiva | SH30079.02 | 10% | 10 mL |
| 2- Mercaptoethanol (1000X) | Gibco | 21985-023 | 1X | 0.1 mL |
| Penicillin-Streptomycin-Glutamine (100X) | Gibco | 10378-016 | 1X | 1 mL |
| Non-Essential Amino Acids (100X) | Gibco | 11140-050 | 0.6X | 0.6 mL |
| HEPES (1M) | Gibco | 15630-080 | 10 mM | 1 mL |
| RPMI Medium 1640 | Gibco | 61870-036 | --- | 37.3 mL |

**Online Table 3.** Reagents used to prepare FACS Buffer. Buffer was kept at 4°C for storage.

| Reagent | Vendor | Identifier | Working concentration |
|---|---|---|---|
| Phosphate Buffered Saline (PBS) | Gibco | 14190-144 | --- |
| Bovine Serum Albumin Solution | SIGMA | A9576 | 0.2% |

**Online Table 4.** Additional reagents for flow cytometry assay.

| Reagent | Vendor | Identifier | Working concentration |
|---|---|---|---|
| Mouse FC Block | BD | 553142 | 1μL/test |
| Brilliant Stain Buffer Plus | BD | 566385 | 10 μL/test |
| Cytofix/Cytoperm™ Plus | BD | 554715 | 100 μL/test of Cytofix, 1X Perm/Wash buffer |
| Sterile Water for Irrigation | Baxter | 2F7114 | ---- |
| Paraformaldehyde 10% Solution | Electron Microscopy Sc. | 15712-S | 200μL of 1% PFA/test |
| Tandem Stabilizer | BioLegend | 421802 | 1:1000 |
| UltraComp eBeads™ Plus | Invitrogen | 01-3333-42 | 1drop/test |
| ArC™ reactive beads | Invitrogen | A10628 A | 1drop/test |
| ArC™ negative beads | Invitrogen | A10628 B | 1drop/test |

**Online Table 5.** Table for flow antibodies used in the optimized 17-parameter panel. APC channel (in green color) was kept blank intentionally to see the effect of the Tandem Stabilizer. CD45 APC was used with reference beads to define the SIM and SSM only. Intracellular cytokines were highlighted by a light gold color. Titer represents the final dilution factors. Each antibody's volume per test (in 100 µL) was also indicated.

| Target | Purpose | Clone | Fluorochrome | Vendor | Identifier | Titer | µL/test | Panel |
|---|---|---|---|---|---|---|---|---|
| CD8 | T cells | 53-6.7 | BUV395 | BD | 563786 | 1:80 | 1.25 µL | Surface |
| CD3 | T cells | 17A2 | BUV496 | BD | 741117 | 1:80 | 1.25 µL | Surface |
| TCR γ/δ | γ/δ T cells | GL3 | BUV563 | BD | 748993 | 1:80 | 1.25 µL | Surface |
| CD62L | Memory | MEL-14 | BUV737 | BD | 612833 | 1:160 | 0.625 µL | Surface |
| IL-4 | Th2 | 11B11 | BV421 | BioLegend | 504119 | 1:80 | 1.25 µL | ICS |
| IL-5 | Th2 | TRFK5 | BV421 | BioLegend | 504311 | 1:160 | 0.625 µL | ICS |
| Viability | | --- | L/D Aqua | Invitrogen | L34966 | 1:500 | --- | Surface |
| CD45R | B cells | RA3-6B2 | BV605 | BioLegend | 103243 | 1:40 | 2.5 µL | Surface |
| NK1.1# | NK cells | PK136 | BV711 | BioLegend | 108745 | 1:80 | 1.25 µL | Surface |
| NKp46# | NK cells | 29A1.4 | BV711 | BioLegend | 137621 | 1:40 | 2.5 µL | Surface |
| IFNγ | Th1 | XMG1.2 | AF488 | BioLegend | 505813 | 1:160 | 0.625 µL | ICS |
| CD44 | Memory | IM7 | Real Blue 744 | BD | 757245 | 1:80 | 1.25 µL | Surface |
| CD45 | Leukocytes | 30-F11 | Real Blue 780 | BD | 569151 | 1:160 | 0.625 µL | Surface |
| IL-2 | Th1 | JES6-5H4 | PE | BioLegend | 503808 | 1:160 | 0.625 µL | ICS |
| CD19 | B Cells | 1D3 | PE-CF594 | BD | 562329 | 1:320 | 0.3125 µL | Surface |
| TNF | Th1 | MP6-XT22 | PE Cy7 | BioLegend | 506324 | 1:160 | 0.625 µL | ICS |
| CD45 | BLANK | 30-F11 | APC | BD | 561018 | 1:320 | 0.3125 µL | BLANK |
| IL-17A | Th17 | TC11-18H10 | R718 | BD | 567098 | 1:80 | 1.25 µL | ICS |
| CD4 | T Cells | RM4-5 | APC Cy7 | BD | 565650 | 1:80 | 1.25 µL | Surface |

#NK1.1 (in C57BL/6) and NKp46 (in BALB/c) were assigned to BV711 and validated using two separate panels.

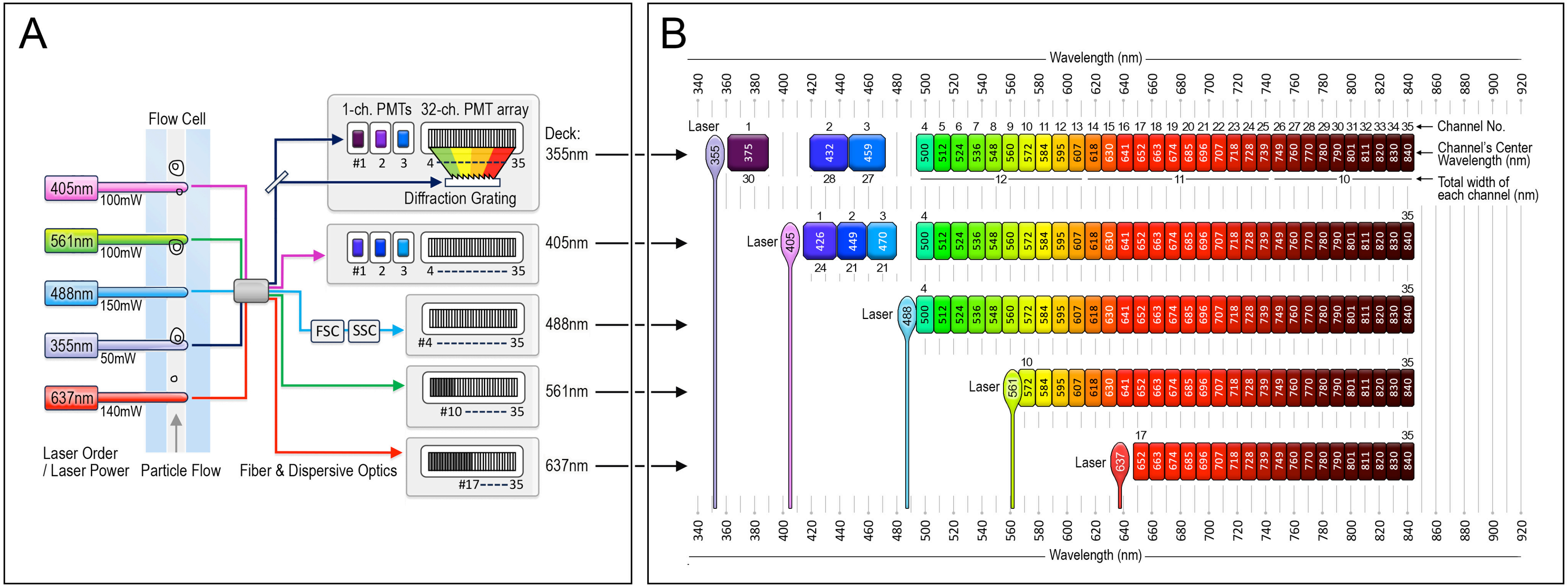


**Online Figure 1:** Graphical representation of the (A) optical layout and (B) configuration of the detectors on the Sony ID7000 spectral cell analyzer. The central wavelength of each detector is given as well as the channel number and total width.

| | BUV395 | BUV496 | BUV563 | BUV737 | BV421 | L/D Aqua | BV605 | BV711 | AF488 | RB744 | RB780 | PE | PE-CF594 | PE-Cy7 | APC | R718 | APC-Cy7 |
|---|---|---|---|---|---|---|---|---|---|---|---|---|---|---|---|---|---|
| BUV395 | 1.00 | | | | | | | | | | | | | | | | |
| BUV496 | 0.02 | 1.00 | | | | | | | | | | | | | | | |
| BUV563 | 0.01 | 0.05 | 1.00 | | | | | | | | | | | | | | |
| BUV737 | 0.01 | 0.00 | 0.01 | 1.00 | | | | | | | | | | | | | |
| BV421 | 0.03 | 0.01 | 0.00 | 0.00 | 1.00 | | | | | | | | | | | | |
| L/D Aqua | 0.00 | 0.64 | 0.06 | 0.00 | 0.01 | 1.00 | | | | | | | | | | | |
| BV605 | 0.00 | 0.00 | 0.12 | 0.01 | 0.00 | 0.00 | 1.00 | | | | | | | | | | |
| BV711 | 0.00 | 0.01 | 0.02 | 0.24 | 0.00 | 0.02 | 0.01 | 1.00 | | | | | | | | | |
| AF488 | 0.00 | 0.00 | 0.00 | 0.00 | 0.00 | 0.00 | 0.01 | 0.01 | 1.00 | | | | | | | | |
| RB744 | 0.00 | 0.00 | 0.01 | 0.03 | 0.00 | 0.01 | 0.01 | 0.00 | 0.00 | 1.00 | | | | | | | |
| RB780 | 0.00 | 0.00 | 0.00 | 0.00 | 0.00 | 0.01 | 0.01 | 0.01 | 0.01 | 0.38 | 1.00 | | | | | | |
| PE | 0.00 | 0.00 | 0.19 | 0.01 | 0.00 | 0.00 | 0.07 | 0.02 | 0.00 | 0.00 | 0.00 | 1.00 | | | | | |
| PE-CF594 | 0.00 | 0.00 | 0.03 | 0.01 | 0.00 | 0.01 | 0.21 | 0.02 | 0.00 | 0.00 | 0.00 | 0.17 | 1.00 | | | | |
| PE-Cy7 | 0.00 | 0.01 | 0.01 | 0.00 | 0.00 | 0.01 | 0.01 | 0.01 | 0.00 | 0.15 | 0.39 | 0.00 | 0.00 | 1.00 | | | |
| APC | 0.00 | 0.00 | 0.01 | 0.00 | 0.00 | 0.01 | 0.00 | 0.01 | 0.00 | 0.00 | 0.01 | 0.00 | 0.00 | 0.01 | 1.00 | | |
| R718 | 0.00 | 0.01 | 0.01 | 0.08 | 0.00 | 0.01 | 0.02 | 0.39 | 0.00 | 0.00 | 0.01 | 0.01 | 0.01 | 0.00 | 0.05 | 1.00 | |
| APC-Cy7 | 0.00 | 0.01 | 0.01 | 0.01 | 0.00 | 0.01 | 0.02 | 0.01 | 0.00 | 0.00 | 0.00 | 0.01 | 0.01 | 0.03 | 0.01 | 0.13 | 1.00 |

**Online Figure 2:** Similarity Index Matrix (SIM) of the optimized 17-color panel on the Sony ID7000 spectral cell analyzer. The SIM was calculated from the spectral references used to unmix the data in this paper. To calculate the similarity score between two fluorophores, normalized fluorescence intensity values of each fluorophore for each detector are plotted and a best fit linear regression line is created. The $r^2$ value of this linear regression line is the similarity score. The similarity score is on a scale of 0-1 and higher scores indicate a higher similarity between two fluorophores.

| Percentile (100 / 99 / 0) | AF488 | APC | APC-Cy7 | BUV395 | BUV496 | BUV563 | BUV737 | BV421 | BV605 | BV711 | L/D Aqua | PE | PE-CF594 | PE-Cy7 | R718 | RB744 | RB780 |
|---|---|---|---|---|---|---|---|---|---|---|---|---|---|---|---|---|---|
| AF488 | 0.00 | 0.00 | 0.00 | 0.00 | 0.38 | 0.16 | 0.07 | 0.00 | 0.07 | 0.00 | 0.19 | 0.06 | 0.15 | 0.00 | 0.00 | 0.41 | 0.26 |
| APC | 0.05 | 0.00 | 0.56 | 0.00 | 0.00 | 0.00 | 0.44 | 0.03 | 0.00 | 0.24 | 0.00 | 0.05 | 0.07 | 0.37 | 1.57 | 0.24 | 0.36 |
| APC-Cy7 | 0.23 | 0.98 | 0.00 | 0.00 | 0.00 | 0.00 | 0.29 | 0.00 | 0.00 | 0.00 | 0.00 | 0.00 | 0.00 | 1.72 | 0.82 | 0.35 | 1.49 |
| BUV395 | 0.03 | 0.00 | 0.00 | 0.00 | 0.64 | 0.18 | 0.09 | 0.03 | 0.03 | 0.03 | 0.11 | 0.09 | 0.00 | 0.00 | 0.05 | 0.02 | 0.00 |
| BUV496 | 0.86 | 0.00 | 0.00 | 1.35 | 0.00 | 1.16 | 0.32 | 0.10 | 0.14 | 0.06 | 1.81 | 0.57 | 0.15 | 0.00 | 0.16 | 0.16 | 0.07 |
| BUV563 | 0.27 | 0.00 | 0.00 | 1.16 | 0.50 | 0.00 | 0.20 | 0.04 | 0.27 | 0.06 | 0.00 | 2.90 | 0.72 | 0.07 | 0.16 | 0.19 | 0.22 |
| BUV737 | 0.14 | 0.16 | 0.96 | 1.31 | 0.49 | 0.21 | 0.00 | 0.11 | 0.00 | 0.37 | 0.10 | 0.10 | 0.00 | 0.44 | 1.69 | 1.73 | 0.93 |
| BV421 | 0.04 | 0.00 | 0.00 | 0.11 | 1.19 | 0.29 | 0.11 | 0.00 | 0.07 | 0.00 | 0.82 | 0.11 | 0.00 | 0.00 | 0.00 | 0.00 | 0.00 |
| BV605 | 0.03 | 0.18 | 0.00 | 0.10 | 0.42 | 0.59 | 0.60 | 0.45 | 0.00 | 0.53 | 0.34 | 1.02 | 1.61 | 0.27 | 0.66 | 0.42 | 0.47 |
| BV711 | 0.14 | 0.43 | 1.43 | 0.64 | 0.60 | 0.19 | 2.66 | 0.77 | 0.00 | 0.00 | 0.44 | 0.03 | 0.00 | 0.72 | 4.59 | 1.44 | 0.92 |
| L/D Aqua | 2.23 | 0.00 | 0.00 | 6.14 | 39.66 | 4.22 | 0.38 | 1.12 | 0.34 | 0.16 | 0.00 | 1.86 | 0.18 | 0.02 | 0.25 | 0.11 | 0.02 |
| PE | 0.19 | 0.00 | 0.00 | 0.00 | 0.04 | 0.38 | 0.07 | 0.01 | 0.28 | 0.00 | 0.00 | 0.00 | 1.01 | 0.15 | 0.08 | 0.25 | 0.35 |
| PE-CF594 | 0.18 | 0.14 | 0.05 | 0.00 | 0.00 | 0.21 | 0.18 | 0.05 | 0.41 | 0.07 | 0.00 | 1.32 | 0.00 | 0.29 | 0.16 | 0.65 | 0.58 |
| PE-Cy7 | 0.64 | 0.15 | 0.57 | 0.00 | 0.00 | 0.13 | 0.23 | 0.00 | 0.12 | 0.00 | 0.00 | 0.84 | 0.34 | 0.00 | 0.22 | 1.07 | 3.31 |
| R718 | 0.00 | 0.00 | 1.66 | 0.00 | 0.00 | 0.00 | 0.89 | 0.08 | 0.00 | 0.40 | 0.00 | 0.00 | 0.00 | 0.75 | 0.00 | 0.39 | 0.65 |
| RB744 | 0.72 | 0.07 | 0.32 | 0.12 | 0.15 | 0.08 | 0.70 | 0.05 | 0.02 | 0.18 | 0.00 | 0.00 | 0.13 | 0.25 | 0.61 | 0.00 | 2.12 |
| RB780 | 1.37 | 0.07 | 0.27 | 0.00 | 0.30 | 0.22 | 0.24 | 0.00 | 0.13 | 0.00 | 0.05 | 0.13 | 0.24 | 0.31 | 0.17 | 1.36 | 0.00 |

**Online Figure 3:** Spillover Spreading Matrix (SSM) of the optimized 17-color panel on the Sony ID7000 spectral cell analyzer. The SSM was calculated from the single-color controls on antibody capture beads (UltraComp eBeads™ Plus) used to calculate the spectral references and used to unmix the data in this paper. Each cell value gives a quantitative assessment of the spillover spreading error that a primary parameter's fluorescence signature (rows/offender fluorophores) creates in the secondary parameter (column, offended fluorophores). Spillover values are presented in magenta color and in percentile format.

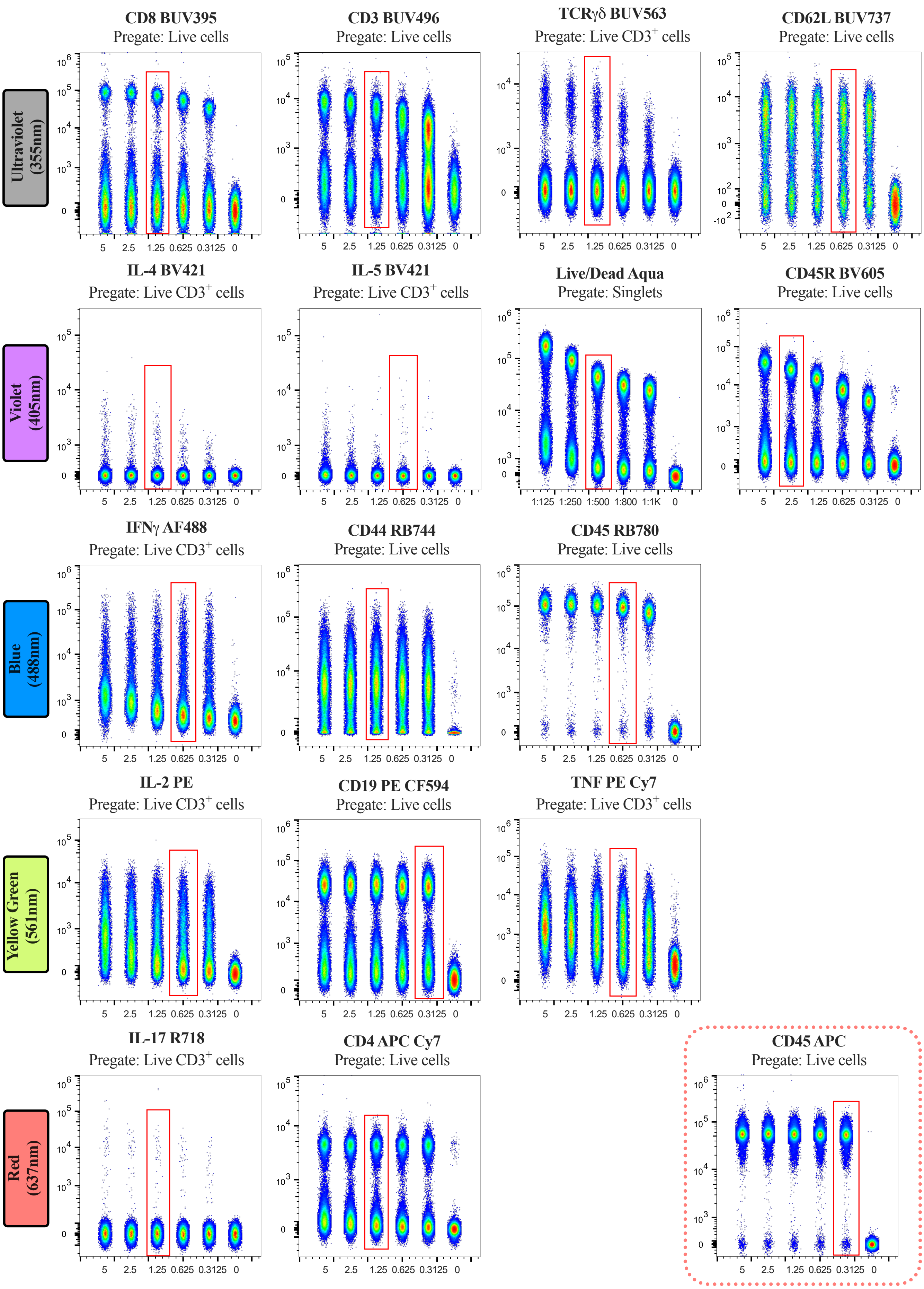


**Online Figure 4:** Optimal concentration of antibodies is defined by titrations.

**Online Figure 4:** (Continued)

Serial fold-dilutions (on the X-axis) of the monoclonal antibodies to detect surface and intracellular cytokines markers were started with 5μL/test. For L/D aqua starting dilution was 1:125. All the surface and intracellular antibodies were titrated using freshly isolated pulled splenocytes from naïve C57BL/6 mice (n=3-5). For intracellular cytokines, cells were stimulated with mitogen along with Brefeldin A to induce cytokine signature and titrated thereafter. FCS files were gated to remove debris (FSC/SSC), doublets followed by gating of live cells or live $CD3^{+}$ cells and concatenated for representation. Red squares portray the chosen titer that provided the distinctive separation between positive and negative populations with minimal background staining. The chosen titration for all the antibodies is also documented as μL/test or as in dilution factor in **Online Table 5**.

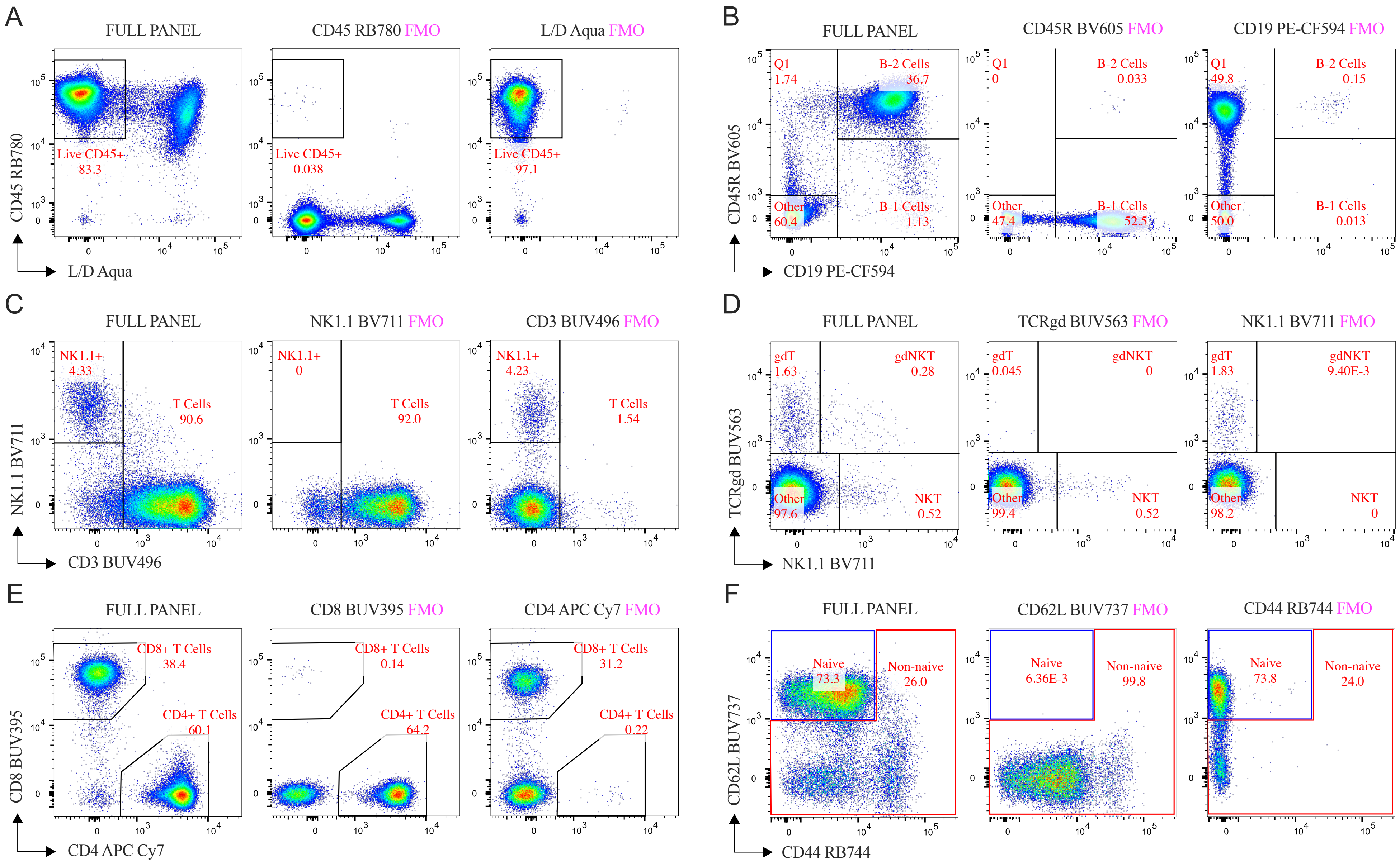


**Online Figure 5:** Defining gating strategy using FMO controls. To establish the accuracy of the manually drawn gates, the following FMO plots are shown for (A) live CD45$^+$ subset, (B) B-1 and B-2 cells, (C) NK1.1 and T cells, (D) γδT, γδNKT and NKT cells, (E) CD4$^+$ and CD8$^+$ T cells and lastly (F) naive and non-naive which is mixed population of effector T cells along with double negative cells. Freshly isolated pulled splenocytes from naïve C57BL/6 mice (n=5) were used here without mitogen stimulation.

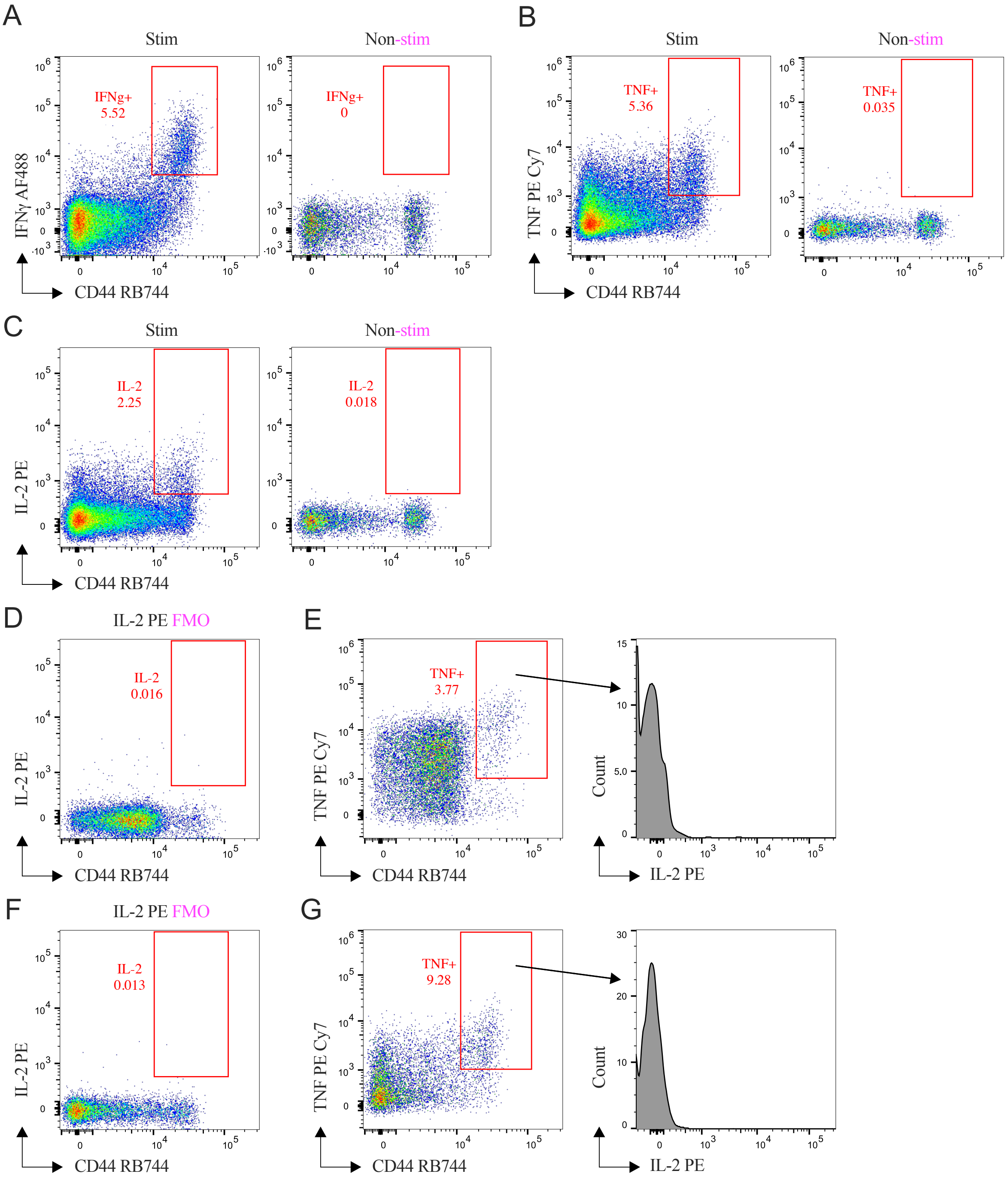


**Online Figure 6:** (A-C) Defining the gating strategy for cytokine$^{+}$ cells in CD8$^{+}$ T cell compartment and (D-F) assessing of PE-Cy7 tandem breakdown. Cytokine signatures (A-C) in stimulated and non-stimulated splenocytes from representative naïve C57BL/6 mice are shown here. (D-G) Freshly isolated pulled splenocytes from naïve C57BL/6 mice (n=5) were used here following mitogen stimulation. (E, G) Full stain with (D, F) PE-FMO control in (D, E) CD4$^{+}$ T cell and (F, G) CD8$^{+}$ T cell compartments were assessed here in the absence of the Tandem Stabilizer. After staining and fixation, cells were preserved without the Tandem Stabilizer and analyzed on Day 1. Absence of any positive signature of TNF PE-Cy7 in PE channel (either in bivariate plot or in histogram) confirmed the absence of PE-Cy7 tandem degradation.

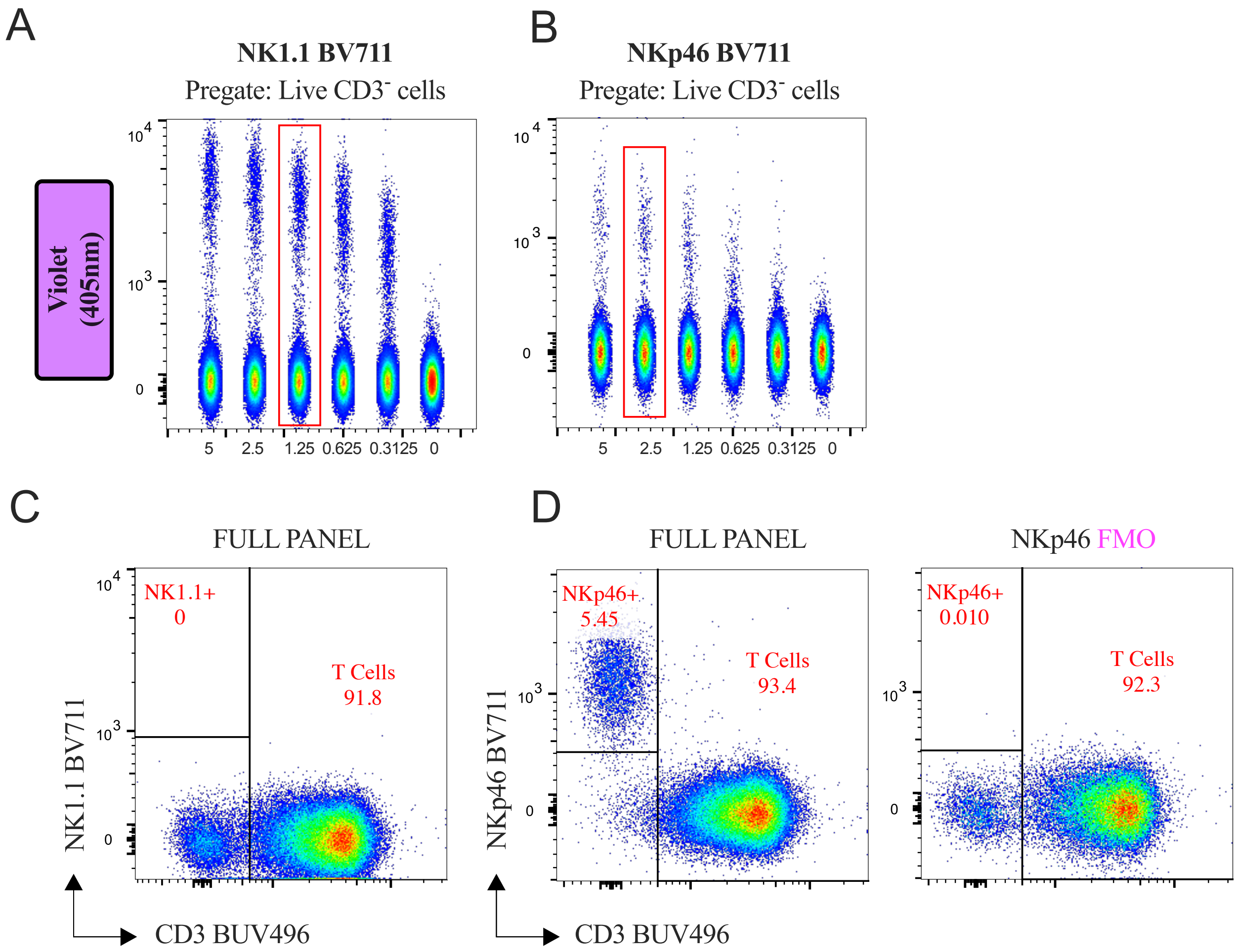


**Online Figure 7:** Marker for NK cells iterations during the panel optimization using C57BL/6 and BALB/c mice. Titrations of (A) NK1.1 BV711 and (B) NKp46 BV711 by serial fold-dilutions (on the X-axis) were started with 5µL/test using pulled splenocytes from (A) naïve C57BL/6 mice and (B) naïve BALB/c mice strains. FCS files were gated to remove debris (FSC/SSC), doublets followed by gating of live CD3$^-$ cells and concatenated for representation. Red squares portray the chosen titer which were also documented as µL/test or as a dilution factor in **Online Table 5**. (C) Incompetency of anti-NK1.1 (PK136) antibody in BALB/c mice splenocytes. (D) Defining the gating strategy using anti-NKp46 (29A1.4) antibody to detect NK cells in BALB/c mice splenocytes by FMO control.

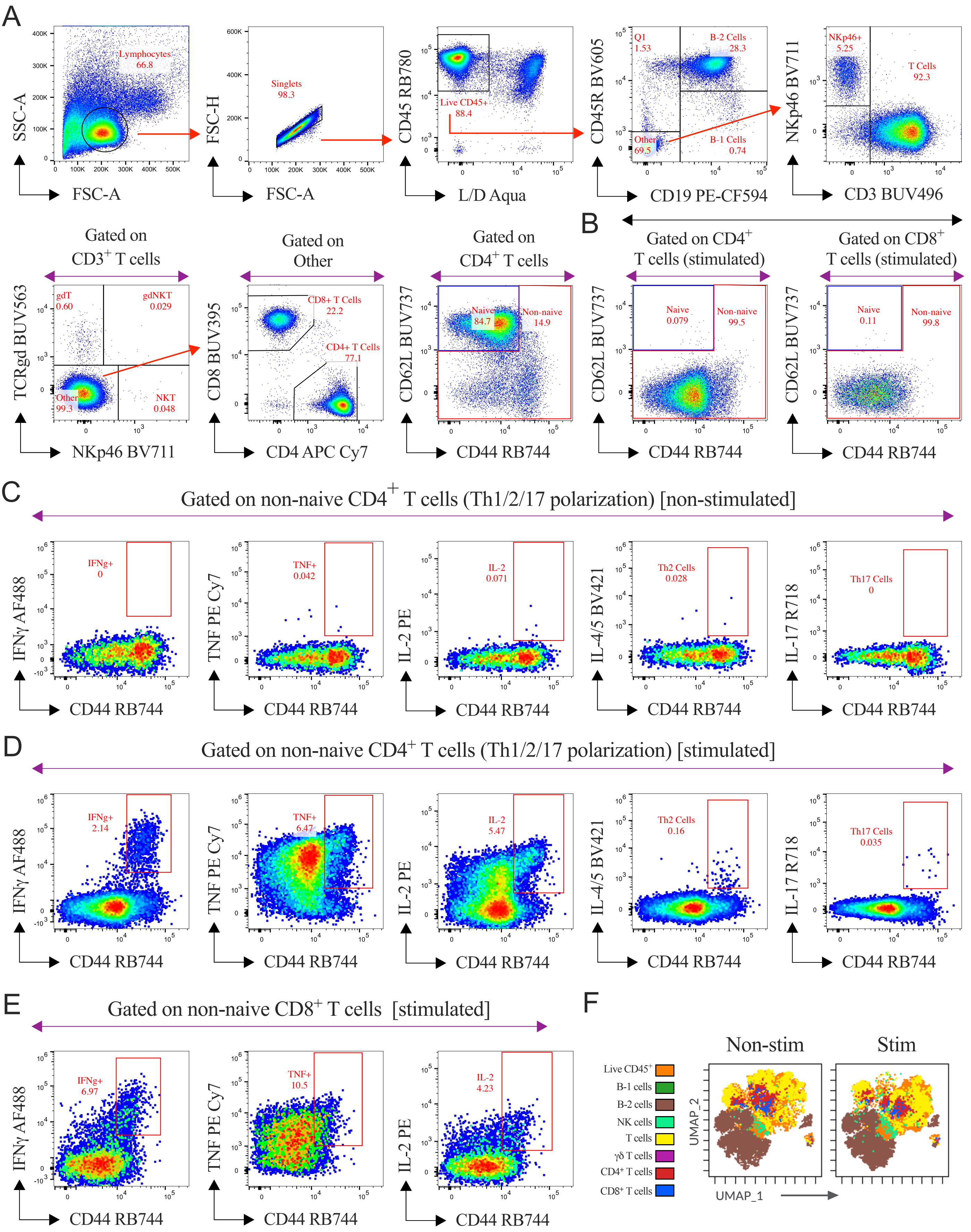


**Online Figure 8:** Panel compatibility in BALB/c mice.

**Online Figure 8:** (Continued)

Representative gating strategy to profile functional cytokines and UMAP analysis for the visualization of the immune cell populations using 17-parameter OMIP-XXX in murine (BALB/c mice) spleen. The rationale of the gating strategy and cell stimulation strategy are mentioned in **Figure 1**.

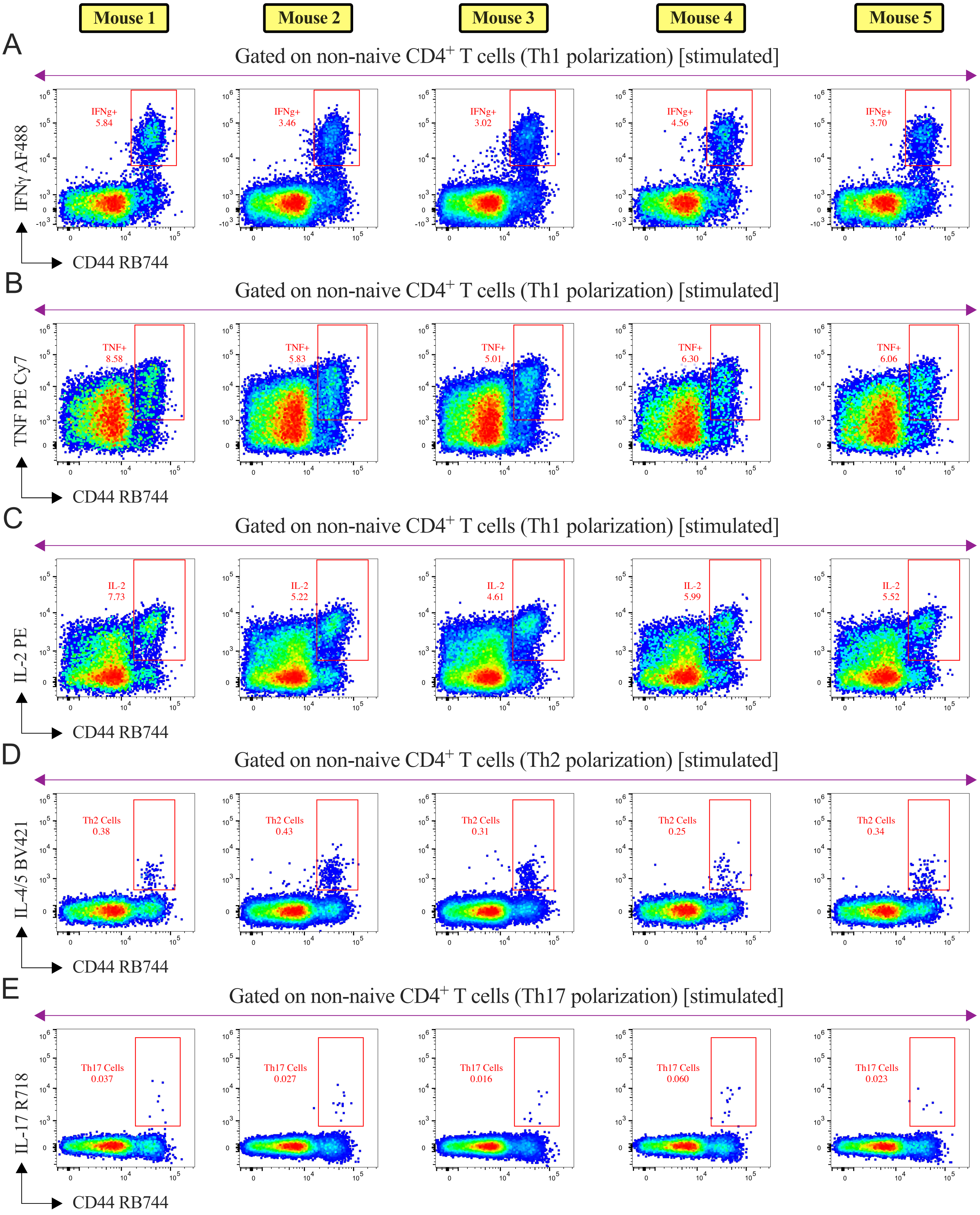


**Online Figure 9:** Consistency of the cytokine signatures (Th1, Th2 & Th17) across C57BL/6 mice strain.

**Online Figure 9:** (Continued)

Freshly isolated splenocytes from C57BL/6 mice strain were stained with the full 16-color (17-parameter) panel after 6h of mitogen stimulation. Five different mice portray (A-C) Th1, (D) Th2 and (E) Th17 signatures in non-naïve (**Figure 1B**) $CD4^{+}$ T cell compartment.

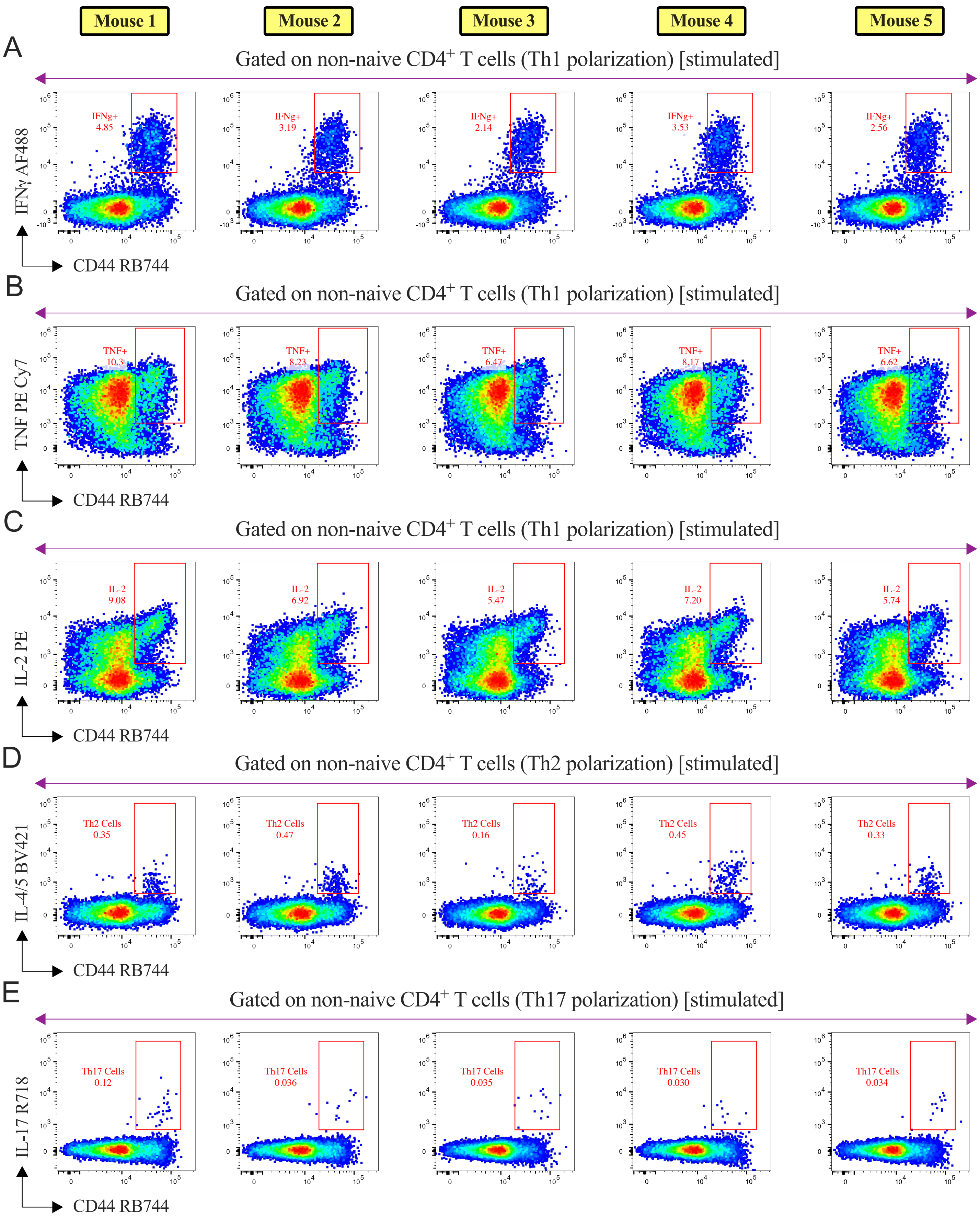


**Online Figure 10:** Consistency of the cytokine signatures (Th1, Th2 & Th17) across BALB/c mice strain.

**Online Figure 10:** (Continued)

Freshly isolated splenocytes from BALB/c mice strain were stained with the full 16-color (17-parameter) panel after 6h of mitogen stimulation. Five different mice portray (A-C) Th1, (D) Th2 and (E) Th17 signatures in non-naïve (**Online Figure 8B**) $CD4^+$ T cell compartment.

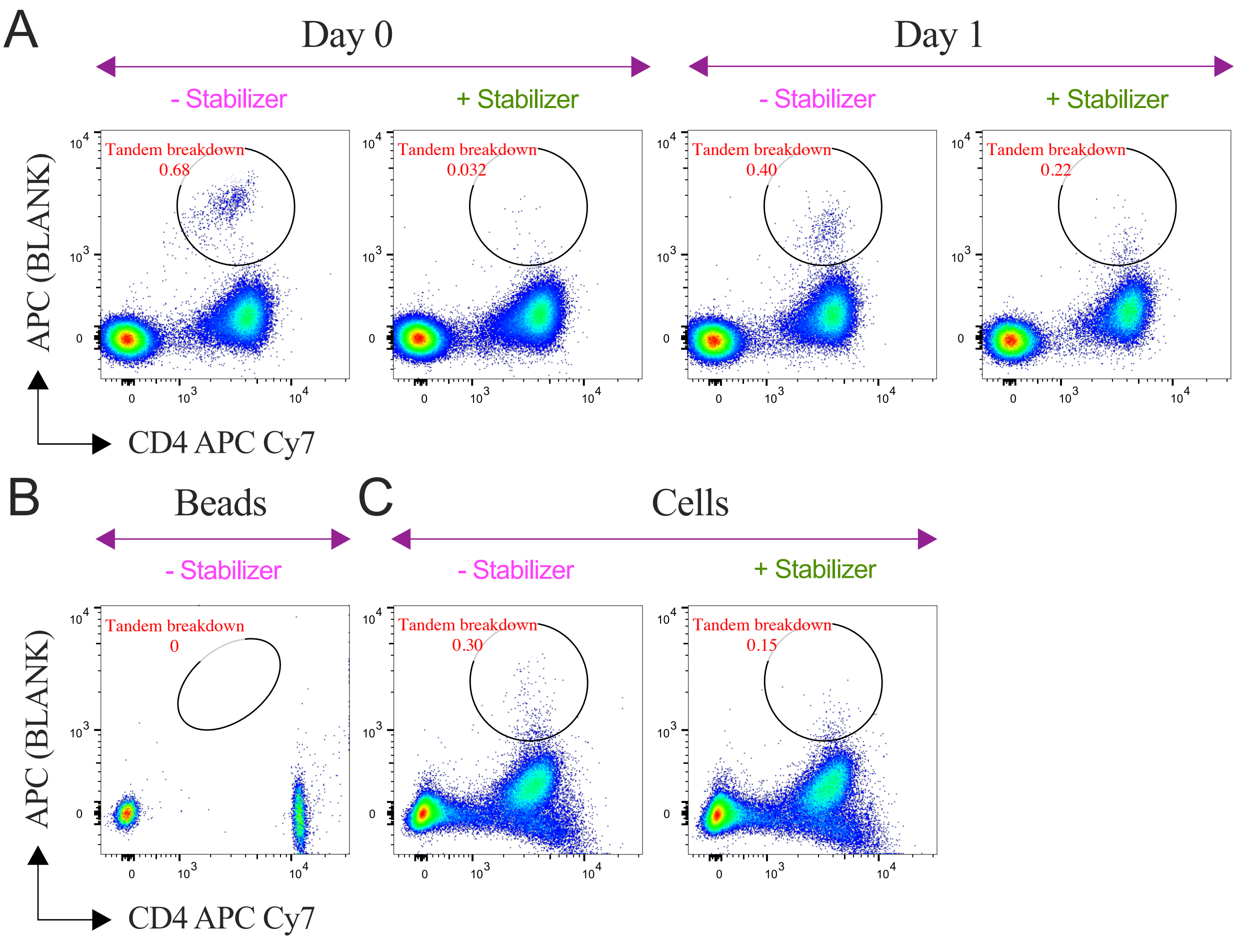


**Online Figure 11:** Reasoning behind using the Tandem Stabilizer. (A) Non-stimulated (Brefeldin A treated) cells were stained with an optimized 16-color panel and left an empty APC channel. After staining and fixation, cells were preserved with or without Tandem Stabilizer (**Online Table 4**) and analyzed in different timepoints (Day 0 and Day 1) to observe the APC Cy7 tandem fluorophore degradation in the blank APC channel. Without the Tandem Stabilizer, the accumulation of tandem breakdown was found to be higher compared with the stabilized group on Day 0 as well as on Day 1. (B) Absence of APC Cy7 tandem fluorophore degradation when using compensation beads (UltraComp eBeads™ Plus). After staining, samples were preserved in PBS and analyzed on Day 1. (C) Single color (APC Cy7) cellular compensation controls were validated by running traditional ICS protocol (mock) using non-stimulated splenocytes. After staining and fixation cells were preserved with or without the Tandem Stabilizer and analyzed on Day 1.

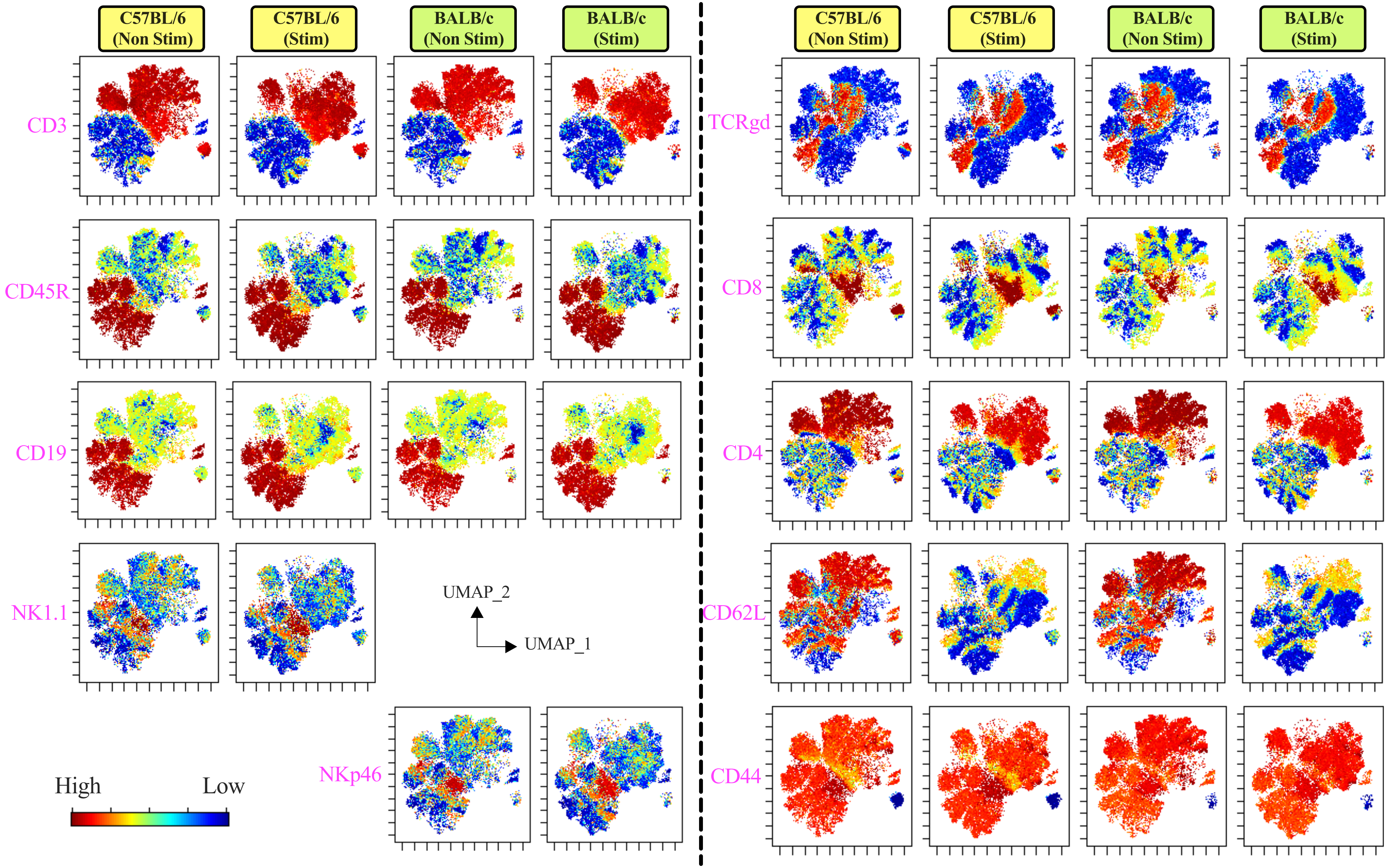


**Online Figure 12:** Dimensionality reduction analysis using the UMAP algorithm and distribution of surface markers in the cellular topography. Surface markers expression (in Z-axis channel) overlaid on UMAP embedding where color indicates median fluorescence intensity of each indicated surface marker expression ranging from low (blue) to high (red). 23K events of live $CD45^{+}$ cells from representative mice were used in dimensionality reduction visualization with UMAP. Splenocytes were either non-stimulated or stimulated with mitogen for 6h.

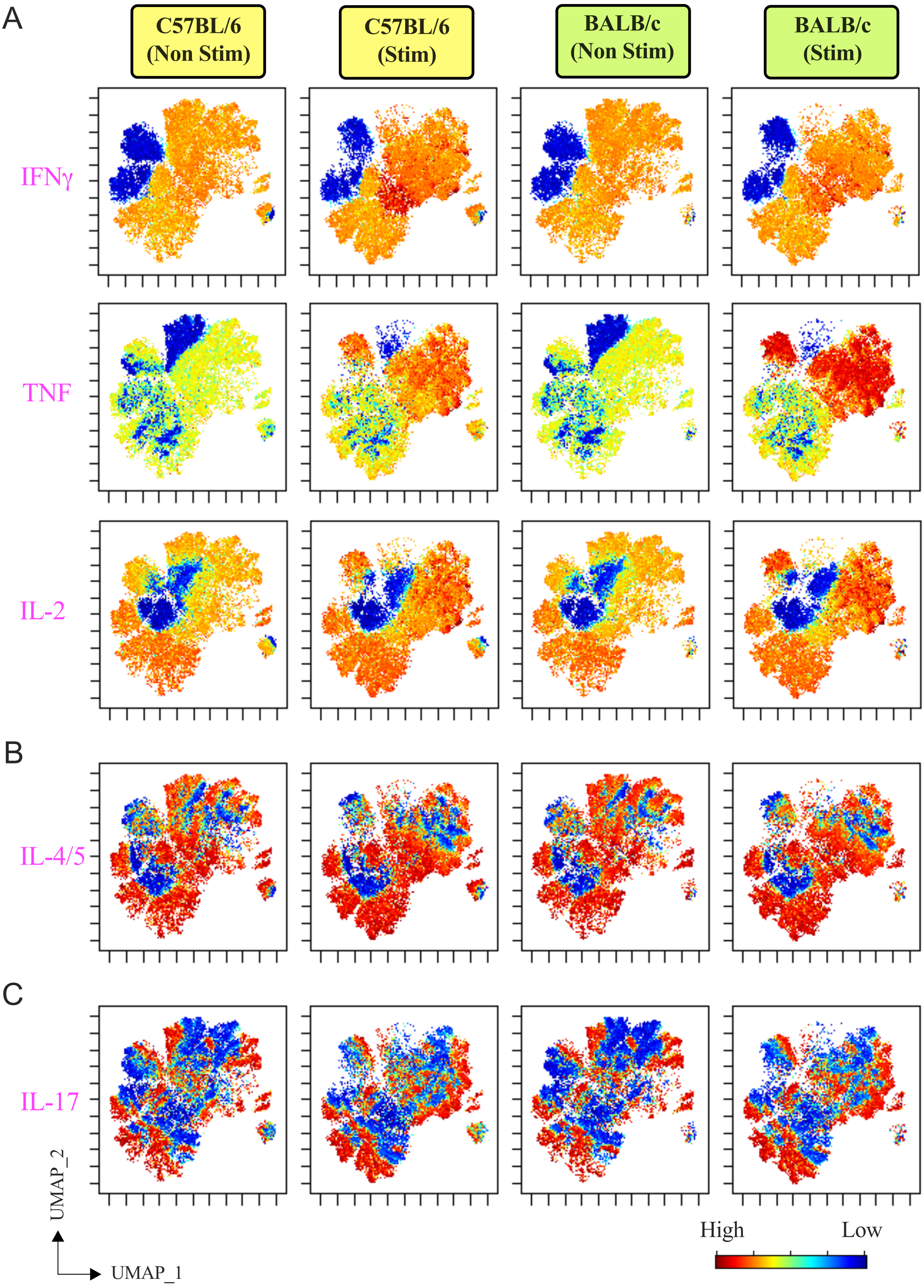


**Online Figure 13:** Dimensionality reduction analysis using UMAP algorithm and distribution of intracellular cytokine markers in the cellular topography.

**Online Figure 13:** (Continued)

Intracellular markers expression (in Z-axis channel) overlaid on UMAP embedding where color indicates median fluorescence intensity of each indicated cytokine marker expression ranging from low (blue) to high (red). 23K events of live $CD45^{+}$ cells from representative mice were used in dimensionality reduction visualization with UMAP. Splenocytes were either non- stimulated or stimulated with mitogen for 6h.
